# The Nature, Extent, and Consequences of Cryptic Genetic Variation in the *opa* Repeats of *Notch* in *Drosophila*

**DOI:** 10.1101/020529

**Authors:** Clinton Rice, Danielle Beekman, Liping Liu, Albert Erives

## Abstract

Polyglutamine (pQ) tracts are abundant in many proteins co-interacting on DNA. The lengths of these pQ tracts can modulate their interaction strengths. However, pQ tracts > 40 residues are pathologically prone to amyloidogenic self-assembly. Here, we assess the extent and consequences of variation in the pQ-encoding *opa* repeats of *Notch* (*N*) in *Drosophila melanogaster*. We use Sanger sequencing to genotype *opa* sequences (5’-CAX repeats), which have resisted assembly using short sequence reads. While the majority of *N* sequences pertain to reference *opa31* (Q_13_HQ_17_) and *opa32* (Q_13_HQ_18_) allelic classes, several rare alleles encode tracts > 32 residues: *opa33a* (Q_14_HQ_18_), *opa33b* (Q_15_HQ_17_), *opa34* (Q_16_HQ_17_), *opa35a1*/*opa35a2* (Q_13_HQ_21_), *opa36* (Q_13_HQ_22_), and *opa37* (Q_13_HQ_23_). Only one rare allele encodes a tract < 31 residues: *opa23* (Q_13_–Q_10_). This *opa23* allele shortens the pQ tract while simultaneously eliminating the interrupting histidine. Homozygotes for the short and long *opa* alleles have defects in sensory bristle organ specification, abdominal patterning, and embryonic survival. Inbred stocks with wild-type *opa31* alleles become more viable when outbred, while an inbred stock with the longer *opa35* becomes less viable after outcrossing to different backgrounds. In contrast, an inbred stock with the short *opa23* allele is semi-viable in both inbred and outbred genetic backgrounds. This *opa23 Notch* allele also produces notched wings when recombined out of the X chromosome. Importantly, *w*^*a*^-linked X balancers carry the *N* allele *opa33b* and suppress *AS-C* insufficiency caused by the *sc*^8^ inversion. Our results demonstrate significant cryptic variation and epistatic sensitivity for the *N* locus, and the need for long read genotyping of key repeat variables underlying gene regulatory networks.

---

Questions about the nature of polyglutamine (pQ) tracts in DNA-binding transcription factors and co-factors have arisen ever since the discovery of the *opa* (5′-CAX) triplet repeats in the *Drosophila* gene encoding *Notch* (*N*) (Wharton *et al.* 1985). This was followed by similar reports of CAG-triplet repeats encoding pQ tracts in other regulatory genes of flies (Kassis *et al.* 1986), yeast (Pinkham *et al.* 1987; Suzuki *et al.* 1988; Bricmont *et al.* 1991; White *et al.* 1991) and other fungi (Yuan *et al.* 1991), mammals (Courey and Tjian 1988), and insect viruses (Carson *et al.* 1991). The significance and interest for human genetic disorders increased after pQ tract expansion was implicated in spinal cerebellar ataxias (SCAs), Huntington’s disease, and other neurodegenerative disorders (Biancalana *et al.* 1992; La Spada *et al.* 1992; Orr *et al.* 1993; Snell *et al.* 1993; Andrew *et al.* 1993).

Study of the Sp1 transcription factor led to the proposal that glutamine-rich regions are transactivation domains associated with co-interacting factors sensitive to *cis*-regulatory binding site spacing (Courey and Tjian 1988; Kadonaga *et al.* 1988; Courey *et al.* 1989). Similarly, it was found that transcriptional enhancers integrating developmental morphogenic signals mediated by pQ-rich factors are sensitive to binding site spacing (Crocker *et al.* 2008, 2010; Crocker and Erives 2013). This led to a series of investigations that strongly implicated the selection of microsatellite repeat (MSR) variants in tuning enhancers targeted by pQ-rich transcription factors (Crocker *et al.* 2008, 2010; Brittain *et al.* 2014). Thus, we suspect that this MSR *cis*-enrichment at enhancers is a consequence of their being targeted by pQ-rich factors, which can also functionally evolve by MSR-related slippage in *trans*. Both *cis*-regulatory and *trans*-regulatory coding variation via MSR variants could affect the degree of pQ *β*-sheet interdigitation between the pQ-tracts of TFs binding to adjacent sites in *cis*. Polyglutamine-rich factors have a tendency to aggregate (Scherzinger *et al.* 1997, 1999; Chen *et al.* 2002), but more specifically, they may do so through the Perutz polar zipper (Perutz *et al.* 1993, 1994). The Perutz polar zipper is a strong *β*-sheet structure created by two adjacent proteins interdigitating together. Furthermore, polyglutamine-rich regions are frequently embedded in intrinsically-disordered domains (Tóth-Petróczy *et al.* 2008), thereby reserving a role for a DNA enhancer scaffold to precipitate complex formation (Brittain *et al.* 2014).

In order to explore the natural relationship between pQ tract length and transcriptional interaction networks, we have focused on the Notch intracellular domain (NICD) for several reasons. First, this domain functions as a mostly dedicated co-activator of the highly-conserved Su(H) transcription factor (Fortini and Artavanis-Tsakonas 1994; Furukawa *et al.* 1995; Lecourtois and Schweisguth 1995; Bailey and Posakony 1995). Second, N acts in a large number of developmental contexts involving tissue-specific enhancers targeted by Su(H) and other pQ-rich transcription factors. Notch pQ variants can thus be assayed through N-target reporter assays and other characterized assays (macrochaete patterning and SOP lineage specification) indicative of Notch activity. Third, NICD contains a single, long pQ tract interrupted only by a single histidine residue. The sequences from both the reference iso-1 (Adams 2000) and Canton-S (Kidd *et al.* 1986) strains feature the wild-type number of 31 consecutive 5′-CAX codon repeats in the eighth exon of *N*, the majority of which are 5′-CAG triplets. We refer to this *opa* repeat configuration of *N* as the *opa31* allelic type. The *opa31* version of *N* encodes the pQ tract, Q_13_HQ_17_. Additional work has shown the existence of a neutral 5′-CAG expansion in some *N* alleles corresponding to the *opa32* type encoding Q_13_HQ_18_ (Tautz 1989; Lyman and Young 1993).

Here we sample the range of *opa* variants in the inbred isofemale lines constituting the *D. melanogaster* genetic reference panel (DGRP), additional classical *N* alleles, X balancers, and other informative stocks. We find an extraordinary range of functionally-variant *N opa* alleles that are invisible to current high-throughput genome sequencing and assembly methods. The distribution of *N opa* length variants is highly asymmetric for *D. melanogaster* and features a long tail of non-wild type alleles in the range from 32–37 residues with unique alleles encoding the intervening histidine at distinct positions within the *opa* repeats. In stark contrast, we found no alleles in the range from 24–30 residues. Remarkably, a single deleterious allele, *opa23*, encodes an uninterrupted pQ tract of 23 residues. This suggests that the histidine plays an important role in attenuating the self-assembly of *β*-sheet secondary structure and that the shortening of the normal pQ tract can be balanced by a simultaneous loss of the histidine. Both short and long extreme *opa* alleles have similar phenotypes, which are suppressed by genomic modifiers present in inbred stocks. We suggest that the *N opa* repeat configuration is an important species-specific, gene regulatory network variable and a major source of post-zygotic species incompatibilities in incipient speciation.

## Materials and Methods

### PCR amplification

Genomic DNA was extracted from small groups and/or single male and/or female flies from each line examined. We used Invitrogen’s Platinum Taq High Fidelity, which is a mixture of: (*i*) recombinant “Platinum” *Taq* DNA polymerase, (*ii*) a *Pyrococcus sp. GB-D* DNA polymerase with 3′ → 5′ exonuclease proofreading activity, and (*iii*) a Platinum *Taq* antibody for hot starts.

#### Amplification of opa repeats

Regions of *Notch* containing the *opa* repeats of exon 8, were amplified via conventional PCR thermocycling (30 cycles of 94*° →T*_*a*_ *→*72*°*) using one of the three following primer pairs and associated annealing temperatures (*T*_*a*_): OPA440f (5′-CAG TCG CGA CCC AGT CTA C) and OPA440r (5′-CCC GGA GAT CCA CAA AAT CCA) with a 58*° T*_*a*_ (D.B. pair used for *N* alleles, X balancers, RAL-100, RAL-105, and wopa23 line), OPA808f (5′-TTA CTT GTT ACA GGC TCG CCA TCG) and OPA808r (5′-CCT CGC TCC AAT CGG AAT TCG) with a 67*° T*_*a*_(C.R. pair used for remaning RAL-lines and *sc* inversions), or OPA720-597f (5′-CCG GCA ATG GAA ATA GCC ACG) and OPA720-597r (5′-AGG GCG GAT TCA TTT GAC CCG) with a 54*° T*_*a*_(C.R. pair used to simultaneously genotype endogenous long 720 bp sequences containing introns 7 and 8, and short 597 bp cDNA transgenes lacking intronic DNA).

#### Amplification of dorsal coding sequence

To amplify and clone coding sequences containing the pQ tract in the Dorsal transcription factor, we used the following primer pair: 5′-GTC GCC ATC GAG CAA CTA C (FWD) and 5′-GCC CGC TAT CGA AGC TAA G (REV).

#### Amplification of In(3R)K-only fragment

To detect the In(3R) Kodani inversion, we used the following primers normally located on the same DNA strand: In(3R)K-R1/F1 (5′-TCG AAG CCCGTG TGG TAA TC), which acts as forward primer if inversion present; and In(3R)K-R3 (5′-TTC TCC CAA CGC ATC ACC AAA). We find that this primer pair amplifies a ∼1,250 bp fragment if the inversion is present.

### Cloning and sequencing

PCR products were purified by gel electrophoresis (1% agarose gel), excision of the band, and gel purification (QIAquick Gel Extraction Kit, Qiagen). Purified PCR products were ligated into the pGEM-T Easy plasmid (Promega) and transformed into JM109 competent *E. coli* (Promega) according to standard protocols. Blue/white screening was used to identify white cells with plasmid containing the insert. Cells from individual white colonies were grown in Luria Broth with ampicillin overnight. Plasmids were isolated from cultures using the QIAPrep Spin Miniprep Kit (Qiagen) and sequenced (ABI BigDye™3.1 and ABI 3730™, Applied Biosystems) with T7, SP6, M13, and/or *N*-specific PCR primers. The RAL lines and *sc* inversion lines were genotyped by sequencing 7–10 indepndent clones with most lines having at least 10 clones and some lines having hundreds of clones sequenced (see Quality Control section of Materials and Methods). The classical *N* alleles, X balancers, the wopa23, RAL-100 and RAL-105 lines were genotyped by sequencing 3–7 clones and many were also confirmed by sequencing a PCR-amplified genomic DNA.

### Quality control experiments for opa genotyping

The commercial *Taq* polymerase + *Psp* polymerase + anti-Taq antibody mixture we used is reported to have 6x higher fidelity than *Taq* DNA polymerase alone. In order to ensure true genotypes could be distinguished from variant clones produced by somatic mutation, PCR mutagenesis, and/or inter- and intra-culture environmental contamination, the following four QC experiments were conducted.

#### QC experiment one

To gauge the extent of PCR sources of error, we conducted eight replicate PCR reactions using as genomic template the miniprep DNA from one *opa31* clone. We obtained sequences for two to three clones from each reaction for a total of 21 sequenced clones. Except for one sequenced clone from a reaction with three sequenced clones all were of the original genotype. The single exception corresponded to an *opa30L* contraction (Q_12_HQ_17_) on the left (L) side of the histidine codon, a PCR error rate of about 4.8%. Similar contraction error rates are seen in our other QC experiments.

#### QC experiment two

To gauge the extent of somatic variation, we used PCR primer pair OPA720-597f/r to clone and sequence 218 long (720 bp) and 276 short (597 bp) *N* clones from BDGP stock #26820, which contains a P-element carrying a *UAS-N-full* cDNA that is presumably non-functional in the absence of a GAL4 driver. Both endogenous and transgenic sequences feature a (Canton-S) wild-type *opa31*3* genotype (shortened to *opa31* here). Of the 218 endogenous clones, 94.0% were of the original *opa31* genotype (205 clones), and 6.0% were contraction variants (nine *opa30L* clones, one *opa29L* clone, and two *opa30R* clones). Of the 276 non-functional cDNA clones that we sequenced, 94.2% were of the original *opa31* genotype (260 clones), and 5.8% were contraction variants (13 *opa30L* clones, two *opa29L* clones, and one *opa27R* clone). Thus, both the functional endogenous locus and the presumably non-functional transgenic loci have identical error rates in a similar range as the PCR-based experiment.

#### QC experiment three

To gauge the extent of intraculture environmental contamination, we sequenced 360 clones from 39 individual larvae and pupae from the RAL-142 stock, which we identified to be polymorphic for *opa31* and *opa32* in equal amounts. We sequenced 5–13 clones per RAL-142 individual with an average of 9.2 clones per individual. Of the 360 clones, 93.6% were of the original genotypes (337 clones of either *opa31* or *opa32*), 5.8% were contraction variants (10 clones each of either *opa30L* or *opa31L*, and one clone of *opa28L*), and 0.6% corresponded to one *opa33L* expansion variant encoding Q_14_HQ_18_ (2 clones). We measure a 144:124 female to male sex ratio in this stock (53.7% female) and thus estimated observing 14 flies for each of *opa31*-only and *opa32*-only genotypes (males and females), and 11 heterozygous genotypes (*opa31*/*opa32* females) under Hardy-Weinberg equilibrium. We observe close to the predicted numbers with 13 *opa31*-only genotypes, 14 *opa32*-only genotypes, and 12 *opa31*/*opa32* heterozygotes if we assume that all individuals with clonal genotype frequencies of one singleton “minor” allele in eight or more sequenced clones are the result of environmental contamination from the DNA of bottle mates of other genotypes. Because there was a statistically impossible number of such true genotype calls, we concluded that these numbers indicated an intraculture environmental contamination rate of 5.7%.

#### QC experiment four

To gauge whether the shortened *opa23* allele in the RAL-646 line has an attenuated or otherwise aberrant mutational rate, we sequenced 358 clones from 38 individual larvae and pupae from this line. These individuals were picked on the same day as the RAL-142 individuals described above using the same picking tool wiped with ethanol between picks. Of the 358 sequenced RAL-646 clones, we identified six and seven clones corresponding to *opa31* and *opa32* alleles, respectively. These correspond to the two RAL-142 alleles, and an interculture contamination rate of 3.6%. These 13 clones were minor contaminants in 10 of the 38 RAL-646 individuals, all of which were found to be homozygous for *opa23*. While this low rate of contamination did not prevent genotyping of any of the 38 individuals, we subsequently adopted more stringent control of reagents and picking tools and eliminated environmental contamination in later *opa* sequencing experiments. QC experiments three and four were conducted at the start of this project, and corresponded to the only time we ever saw evidence of environmental contamination.

Putting aside the 13 *opa31*/*opa32* contaminant clones, we are left with 340 total clones sequenced from this stock. Of these, 96.5% were of the original *opa23* genotype (333 clones), and 3.5% were *opa22L* contractions (12). In summary, the contraction variants were identical to the (CAG)_7_ *→* (CAG)_6_ contractions seen in all the other QC experiments, including the PCR control.

### DGRP line outcrossing

To produce our outcrossed lines, 1–5 virgin RAL females (from RAL-# parent stock lines) were crossed with 2–3 X balancer males from lab stocks of *FM7c/N*^1^ (background one) or *FM7a* (background two). We performed five generations of outcrossing of virgin females to the balancer stocks males, followed by two generations of inbreeding to re-homozygose the RAL X chromosome. We refer to these lines as the RALX#-bg1 and RALX#-bg2 outcrossed lines.

To produce the *w*^1118^ *N*^*opa*23^ line (“wopa23”), we first crossed RAL-646 (*opa23*) virgin females with *w*^1118^ males (P_0_). Second, we took heterozygous F_1_ virgin females and crossed them to FM7c males. Based on the distance between *w* and *N* and published recombination rates for this region, we expected a recombination rate of 1% (Comeron *et al.* 2012). So, third, we took 202 individual white-eyed F_2_ males, choosing as much as possible those that might have ectopic macrochaetes, and set up individual crosses with FM7a virgin females. Fourth, after F_3_ larvae were visible, we then genotyped the *opa* repeats from the single F_2_ males by Sanger sequencing of amplified genomic DNA. Fifth, after about 50 genotypes, we genotyped our first white-eyed *opa23* male from one of the F_2_ crosses. We took the F_3_ virgin females and crossed them to FM7a sibling males. Last, we crossed the non-Bar, white-eyed F_4_ males to heterozygous sibling females, and selected only white eyed non-balancer flies in the F_5_ generation to homozygose the recombined *w opa23* X chromosome.

### Embryonic survival assays

Populations of 400–600 parent flies from individual DGRP parent stocks (RAL-# lines) or outcrossed lines (RALX#-bg1 and RALX#-bg2) were raised at room temperature (22–24*°*) and allowed to lay eggs on a blue-dyed apple juice agar plate at 25*°* for four hours 1–12 days after eclosion. Efforts were made to standardize the size of the parent population, as well as the age spectra. Embryos were counted in sections of the apple juice agar plates so that similar numbers and densities of embryos were assessed across lines. These sections were placed in the middle of larger apple juice (AJ) agar plates with yeast paste smeared around the outer portion of the plate to attract larvae away from the middle. The fraction of embryos failing to hatch was determined by counting the number of unhatched embryos approximately 40 hours after the end of the laying period.

### Macrochaete scoring

The ectopic macrochaete phenotypic data shown in the Fig. 4 graphs involved scoring doubled (non-split) anterior and posterior dorsocentral macrochaetes (DCs), anterior and posterior scutellar macrochaetes (SCs), and posterior post-alars (pPAs).

The macrochaete frequency plots for the 13 macrochaetes of the hemi-notum involved the following scoring system applied to each of the 13 macrochaete locations. A score of 1.0 was assigned for a normal looking macrochaete (normal thick bristle) at a correct location. A score of 0.0 was assigned for a missing bristle, and the presence of an empty socket was recorded if observed. A score of 2.0 was assigned if an ectopic bristle was observed near a normal position, or if a split bristle was observed (two bristles emanating from a single socket). Split bristles were rare and such cases were recorded if observed. A score of 0.5 or other fractional values were assigned for abnormally slender bristles that were either larger than a microchaete or emanating from a robust macrochaete-type socket. Occasionally, an ectopic bristle was observed exactly mid-distance from the anterior and posterior dorsocentrals or scutellars, and in such cases each bristle position was assigned a value of 1.5. A score of –1.0 (1.0*–*2 defects = –1.0) was assigned when an empty or missing socket at a correct location was observed together with a nearby ectopic macrochaete. Rarely, a score of –2.0 (1.0 –3 defects = –2.0) was assigned when two empty sockets replaced a single normal macrochaete. Rarely, the data for a specific bristle position for a given genotype involved *both* ectopic (> 1.0 scores) and missing bristles (< 1.0 scores). To prevent the cancellation or masking of such defects using the described scoring system, only the more frequent defect (ectopic or missing bristle but not both) was scored, with the data for the less frequent defect being assigned 1’s. Typically, 40–60 hemi-nota from 20–30 adult flies of each sex were scored, but more were scored when many of the flies had wild-type macrochaete patterning.

### Western immunoblots

Embryonic lysates were obtained from timed egg collections on AJ plates and aged to three different time points: *t*_1_ (0–2 h), *t*_2_ (2–4 h), and *t*_3_ (4–6 h). SDS-PAGE (BIO-RAD TGX Stain-Free™10% FastCast acrylamide) gels were loaded with 10 or 15 *μ*L of protein extracts in loading buffer and typically run for 1 h at 128 V using BIO-RAD casting system and electrophoresis apparatus (Mini-PROTEAN Tetra Cell). Gel-separated proteins were transferred to PVDF blotting paper and blocked overnight at 4*°* with 5% non-fat milk in PBT (1x PBS + 0.1% tween-80). Blots were then incubated with one or two of the primary antibodies diluted 1:100 in PBT-1.0 (1X PBS + 1% Tween-80) for 1 hour shaking at room temperature (RT). Mouse C17.9C6 (Fehon *et al.* 1990) and E7 monoclonal antibodies, which are specific for *Drosophila* Notch intracellular domain and *β*-tubulin respectively, were obtained from the Developmental Studies Hybridoma Bank (U. of Iowa) and used as primary antibodies. After 4x washes in 1x PBT (15 minutes per wash), the blots were incubated 1 hr at RT shaking with HRP-conjugated goat anti-mouse secondary antibodies at a 1:500 dilution. PVDF paper was washed 4x in PBT (15 minutes per wash) and marked to indicate the location of the rainbow protein marker bands before being incubated with ECL detection solutions per manufacturer’s instructions (Thermo Scientific SuperSignal West Chemiluminescent substrate, or Bio-Rad Clarity Western ECL substrate). The blot paper was then inserted in a plastic envelope and exposed to film.

## Results

### 1. Naming the many opa variations

The *opa* nucleotide repeats are located in the eighth exon of *Notch* (*N*) and encode two polyglutamine (pQ) tracts separated by a single histidine (Fig. 1). This pQ-H-pQ sequence is part of the N intracellular domain (NICD), which is translocated into the nucleus upon cleavage (Fig. 1A). These pQ tracts are intrinsically disordered and surrounded by many other similar peaks of disorder (Fig. 1B). While the NICD pQ tracts are immediately flanked by conserved amino acid sequence, we find evidence of multiple *opa* repeat configurations that are unique to the *N* gene of *D. melanogaster* (Fig. 1C). To better relate our phenotypic results from studying variant *N opa* alleles, we introduce a simple nomenclature for the different *opa*-encoded pQ configurations.

**Figure 1.**
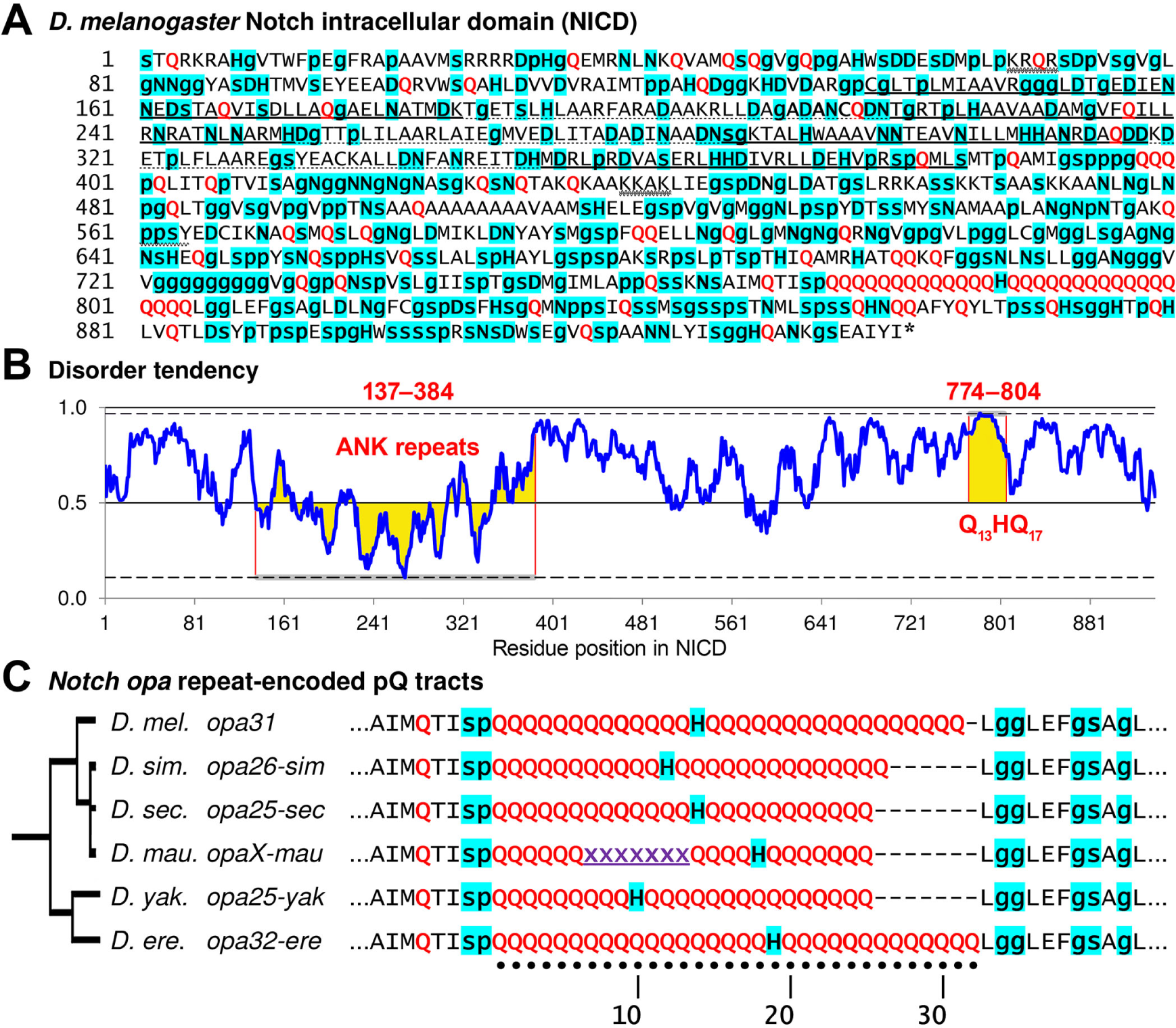
The cleaved Notch intracellular domain (NICD) is characterized by a polyglutamine (pQ) tract configuration that is unique to each *Drosophila* species. **(A)** Shown is the 939 residue long polypeptide sequence for NICD from *D. melanogaster*. Residues that tend to be secondary structure breakers are highlighted in cyan (p, g, s, D, N, and H) with the smaller amino acids shown in lower-case lettering. Glutamines (Qs) are shown in red. The seven ankyrin repeats are shown in alternating bold and dotted underlining. Two nuclear localization sequences, NLS1 (KRQR) and NLS2 (KKAK), are indicated with double wavy underlining. The Nedd4 ubiquitination site (ppsY) is also indicated in light wavy underlining. **(B)** Much of the NICD is disordered as demonstrated by this plot of long disorder tendency based on pairwise energy content (IUPred) (Dosztányi *et al.* 2005a,b). The ankyrin (ANK) repeats (residue positions 137–384) and the pQ tracts (Q_13_HQ_17_ at residue positions 774–804) are notable for having the least and most disorder tendencies, respectively (highlighted in yellow). **(C)** The *opa*-encoded pQ repeats of *Drosophila* are characterized by two adjacent pQ tracts separated by a single conserved histidine. While the surrounding amino acids are conserved, each species features a unique pQ configuration characterized by the length of the pQ tracts on either side of the histidine. The assembly for *D. mauritiana* indicates a unique position for the His residue even though the entire *opa* repeat sequence is still uncertain (Garrigan *et al.* 2012). In this study, we refer to different *N opa* configurations by appending the length suffix to the word “opa”. We add an additional index when more than one configuration exists for a given length as distinguished by the position of the single histidine (*e.g.*, *opa33a* and *opa33b*). Occasionally, we add an additional number when two independently-derived nucleotide sequences encode the same pQ tract configuration (*e.g.*, *opa35a1* and *opa35a2*; see Figs. 2 and 3). Unless a species is indicated (*e.g.*, *opa26-sim*), all *opa* designations refer to alleles from *D. melanogaster*.

The single intervening histidine (His) is relevant to our findings and is a convenient marker for naming distinct *opa* configurations even when they share the same number of repeats. The His codon (5′-CAY) is closely related to the codon for glutamine (Gln) (5′-CAR) and so one might expect to frequently observe His codon turnover within this tract, but this is not the case. The His codon changes position only indirectly after changes in the lengths of the two flanking pQ tracts. Thus, it is likely that the single His residue is highly conserved for a specific role. For example, His residues are known *β*-sheet breakers and have been found to attenuate the *β*-sheet forming potential of pQ peptides (Sharma *et al.* 1999; Sen *et al.* 2003; Kim 2013). Interestingly, species that are closely related to *D. melanogaster* each possess a single His residue at a unique position within the pQ tract (Fig. 1C).

To first distinguish the variants by repeat length, we refer to each *N opa* configuration by a length designation. For example, here *opa31* will refer to the *opa* nucleotide repeats encoding the 31 residues Q_13_HQ_17_. Remarkably given subsequent findings to be described, we have never observed an *opa31* variant allele for *D. melanogaster* that possesses a shifted His residue while maintaining a length of 31 repeats. Nonetheless, we had to use an additional lower-case suffix letter for cases in which there were two or more observed allelic classes of the same length. For example, here *opa33a* will designate Q_14_HQ_18_, *opa33b* will designate Q_15_HQ_17_, and *opa33c* will designate Q_13_HQ_19_.

Occasionally, some identical Q_*n*_HQ_*m*_ configurations are encoded by independently-derived *opa* variant nucleotide sequences. This is possible to determine because of the unique patterns of 5′-CAG and 5′-CAA codons. When we find identical pQ configurations encoded by independently-derived nucleotide sequences, we keep the same letter designation but append an additional number to the letter suffix. Thus, for example, *opa35a1* and *opa35a2* both encode Q_13_HQ_21_ but have been derived by independent histories of insertions and deletions (see Fig. 3).

**Figure 2.**
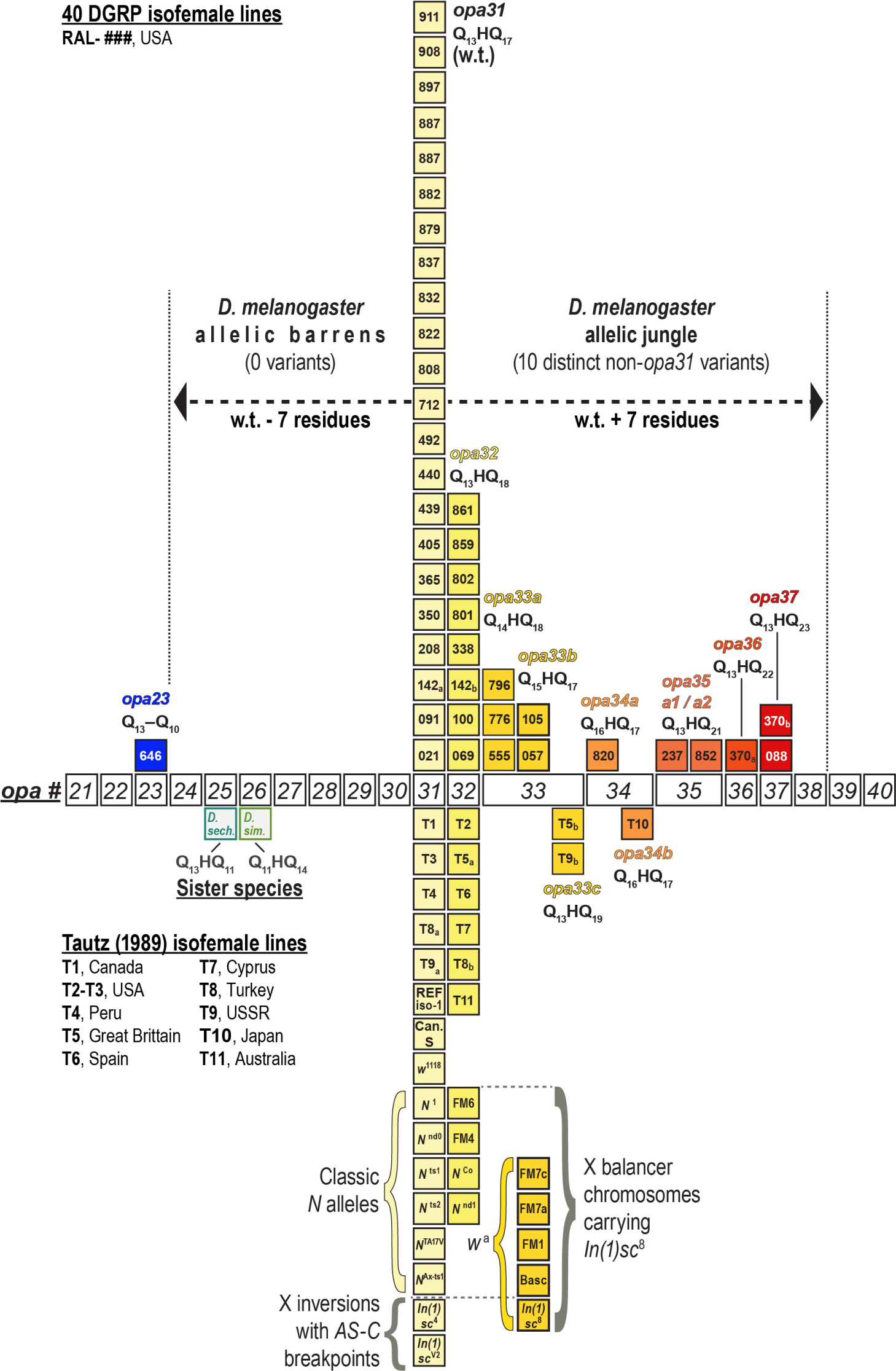
Shown are the *opa* genotypes of 40 DGRP lines (numbered boxes above line), which we genotyped by Sanger sequencing of multiple independent clones. Genotypes are shown as a distribution of *opa* variants from short (left, bluer boxes) to long (right, redder boxes) with each column representing a distinct *opa* allele. Homozygous genotypes and heterozygous haplotypes (142a/142b and 370a/370b) are each represented by a single box. The distribution of *opa* alleles from *D. melanogaster* is highly asymmetric and does not harbor any alleles encoding < 31 residues except for *opa23*, which is much shorter and missing the His codon. Because of this asymmetry, we refer to the range *opa30*–*opa24* as the *D. melanogaster* allelic barrens and the range *opa32*–*opa38* as the allelic jungle. Below the *opa-#* axis are shown several classic *N* alleles, X balancers, and X inversions which we also genotyped for various reasons explained in the text. Canton-S wild-type and *N*^*Co*^ and *N*^*nd*^ mutant alleles have been previously reported (Kidd *et al.* 1986; Lyman and Young 1993). Also shown is the distribution of *opa* genotypes for 11 isofemale lines from different world-wide populations (Tautz 1989). This smaller distribution conforms to the RAL distribution and adds two different rare variants to the allelic jungle (*opa33c* and *opa34b*).

**Figure 3.**
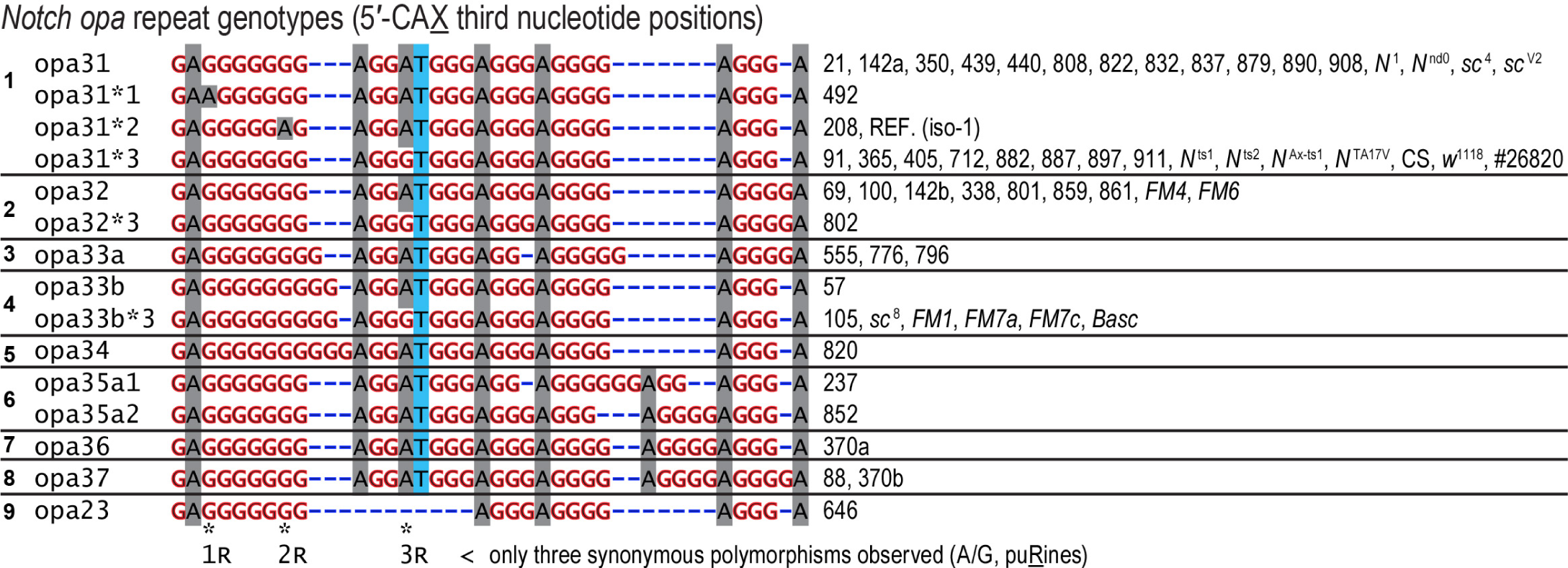
Diverse RAL nucleotide sequences for the *opa* repeats of *D. melanogaster* encode 9 distinct pQ-isoforms of Notch. The *opa* repeat genotypes (third codon positions of 5′-CAX shown) also demonstrate that the longest repeats of 5′-CAG coincide with the most polymorphic regions, consistent with the effect of repeat length on replication slippage frequency and nucleotide substitution rates. We also note that the 3’ end of the *opa* repeats corresponds to a repeat of repeats, (GGGGA)_2–3_, and derivatives thereof. This is also a hypervariable site. Canton-S (CS), BDGP stock #26820 used in QC experiments, and other genotyped stocks are also listed.

The most frequent *opa* alleles, wild-type *opa31* and *opa32*, also have the most derived variants caused by single synonymous substitutions at the nucleotide sequence level. We will usually not distinguish between these and refer to them non-specifically with an asterisk (*opa31\**) only when discussing features at the nucleotide level. (In this study we noted three such polymorphisms and refer to them by applying the suffixes *1, *2, and *3 to the *opa* genotype base name).

#### Quality control experiments

We first conducted an extensive set of quality control (QC) sequencing experiments using genomic DNA amplified from homozygous and heterozygous RAL individuals and clonal mini-prep DNA of defined genotypes (see Materials and Methods). These QC methods ensured that we could reliably and consistently genotype these lines and rule out sources of error from PCR mutagenesis, somatic mutation, inter- and intra-culture contamination, and PCR-crossover effects in cases of mixed *opa* variant DNAs in one sample. We find that there is a small but measurable rate of contraction error that goes from a low of 3.5% in the RAL-646 *opa23* stock to a high of 5.8% and 6.0% from a stock carrying a non-functional full-length *N* cDNA transgene (*UAS:N*^*FL*^) and endogenous locus, respectively (BDGP stock #26820). RAL-142, one of the few RAL-lines heterozygous for *opa* variants, has a similar 5.8% contraction rate, but also was found to have expansion variants (the same two *opa33L* clones in 360 sequenced clones) for an overall error rate of 6.4%, which nonetheless is insufficient to prevent accurate genotyping of any single individual fly. A smaller 4.8% contraction rate in sequenced clones was seen when we amplified from a mini-prep DNA plasmid clone of *opa31* genotype. Thus, in summary, the overwhelming majority of non-genotypic variant clones, which we sequenced at a rate of 3–6%, predominantly originate from PCR-based contractions. These clones do not correspond to variants seen in the wild (compare to the *opa* “allelic barrens” described below) and they do not overwhelm our ability to genotype single individuals.

We typically attempted to sequence ten independent clones from each genomic prep, but in some cases we sequenced a PCR amplification followed by sequencing of 3–7 sub-clones for confirmation. Last, we also re-genotyped many lines and strains after several months without getting different genotypes.

### 2. N opa variants from the DGRP RAL lines

To sample the natural distribution of *N opa* alleles, we made use of the 205 inbred isofemale lines constituting the *Drosophila melanogaster* Genetic Reference Panel (DGRP) (Mackay *et al.* 2012). The DGRP stocks were founded by 1,500 mated females of *D. melanogaster* caught in 2003 outside of a farmers market in Raleigh (RAL), North Carolina, and individually subjected to 20 generations of full-sibling inbreeding (Mackay *et al.* 2012). The initial bulk set of DGRP genome assemblies were based on paired short reads (Illumina 75 nucleotides), which are incapable of individually spanning the 93 base pairs constituting the typical *N opa* sequence and resolving alignment of its repeat structures. These were subsequently complemented with 454 reads as well as various consensus assembly strategies (Mackay *et al.* 2012; Huang *et al.* 2014). The DGRP Freeze 1.0 assemblies focused on identifying high-quality SNPs with significant minor allele frequencies (Huang *et al.* 2014). Subsequent re-assemblies were used to establish an assembly consensus in order to improve the sequence genotyping around micro-satellite repeat structures (Huang *et al.* 2014). However, in our experience with Sanger re-sequencing of the *N opa* repeats, the DGRP 1.0 and similar DGRP 2.0 assemblies are unreliable at this locus (Table 1). For example, of the 158 pre-freeze PopDrowser assemblies based only on Illumina reads (Ràmia *et al.* 2012), over 40% of *opa* sequences contain N’s (ambiguous base calls) and/or 5′-CAR codons. Similarly, of the 162 DGRP Freeze 1.0 assemblies based only on Illumina reads and aligned using different methods (Mackay *et al.* 2012), 33% of *opa* sequences contain N’s.

**Table 1.** Sampling the cryptic Notch opa genotype via Sanger sequencing

| Group | % non-*opa31*<sup>a</sup> | Avg. length deviation<sup>b</sup> | RAL lines (N *opa* genotype) |
| --- | --- | --- | --- |
| N Dirty Dozen<sup>c</sup> | 58.3% (7/12) | 1.67 a.a. | 21 (*opa31*), 237 (*opa35a1*), 350 (*opa31*), 439 (*opa31*), 440 (*opa31*), 555 (*opa33a*), 646 (*opa23*), 776 (*opa33a*), 796 (*opa33a*), 801 (*opa32*), 802 (*opa32*), 822 (*opa31*) |
| Radiation Resistant<sup>d</sup> | 61.8% (10.5/17) | 1.94 a.a. | 91 (*opa31*), 69 (*opa32*), 338 (*opa32*), 208 (*opa31*), 57 (*opa33b*), 492 (*opa31*), 142 (*opa31/32*), 879 (*opa31*), 370 (*opa36/opa37*), 808 (*opa31*), 237 (*opa35a1*), 88 (*opa37*), 405 (*opa31*), 646 (*opa23*), 801 (*opa32*), 105 (*opa33b*), 776 (*opa33a*) |
| Radiation Sensitive<sup>e</sup> | 28.6% (6/21) | 0.57 a.a. | 21 (*opa31*), 350 (*opa31*), 365 (*opa31*), 439 (*opa31*), 440 (*opa31*), 712 (*opa31*), 796 (*opa33a*), 802 (*opa32*), 820 (*opa34*), 822 (*opa31*), 832 (*opa31*), 837 (*opa31*), 852 (*opa35a2*), 859 (*opa32*), 861 (*opa32*), 882 (*opa31*), 887 (*opa31*), 890 (*opa31*), 897 (*opa31*), 908 (*opa31*), 911 (*opa31*) |
| *In(3R)K/In(3R)K*<sup>f</sup> | 100.0% (3/3) | 3.67 a.a. | 100 (*opa32*), 105 (*opa33b*), 646 (*opa23*) |

<sup>a</sup> Each haplotype in the heterozygous RAL-142 and RAL-370 stocks counted as 0.5.
<sup>b</sup> Average absolute value of deviation from 31 amino acids (a.a.)
<sup>c</sup> DGRP1.0 lines with ambiguous assembly sequence (contains N's) or His → Gln (RAL-646)
<sup>d</sup> Top radiation-resistant RAL lines genotyped, in order of avg. survival (Vaisnav *et al.* 2014): 91 (49.0/50 flies), 69 (47.5/50 flies), 338 (45.0/50 flies), 208 (44.5/50 flies), 57 (43.0/50 flies), 492 (42.0/50 flies), 142 (41.0/50 flies), 879 (39.0/50 flies), 370 (49/50 flies), 808 (31.5/50 flies), 237 (28.5/50 flies), 88 (26.5/50 flies), 405 (26.0/50 flies), 646 (9.5/50 flies), 801 (4.5/50 flies), 105 (2.0/50 flies), 776 (2.0/50 flies)
<sup>e</sup> Radiation-sensitive RAL lines genotyped [0/50 flies surviving (Vaisnav *et al.* 2014)]
<sup>f</sup> Only three RAL lines are homozygous for Kodani inversion (Huang *et al.* 2014).

The difficulty of aligning lengthy repeat structures is compounded by the alternate sub-alignments made possible by the repeat structure (Huang *et al.* 2014). This is further exacerbated at the *N opa* region by the strategy of mapping read alignments to the reference sequence (iso-1, an isogenic strain with genotype *y*; *cn bw sp*), which contains a minor synonymous polymorphism (5′-CAG*→* 5′-CAA) (Brizuela *et al.* 1994; Adams 2000).

Figure 2 (distribution of pQ variants) and Figure 3 (nucleotide sequences) show the results of our Sanger-sequencing of several groups of DGRP lines, which we chose by various strategies listed in Table 1. First, after consulting the polished DGRP1.0 and DGRP2.0 sequences, we had a prior expectation of finding few length variants different from *opa31*. We therefore chose to sequence eleven lines, which had N’s annotated in their *N opa* assemblies, and one line (RAL-646), which was annotated as a simple coding substitution changing the central histidine (H) to glutamine (Q). We refer to these as the Notch Dirty Dozen, and this sampling of the RAL lines resulted in the identification of three uncommon and previously unreported alleles: *opa23* (encoding Q_23_), *opa33a* (encoding Q_14_HQ_18_), and *opa35a1* (encoding Q_13_HQ_21_) (Fig. 2, Table 1).

The Notch Dirty Dozen was not a random sampling of the DGRP stocks (RAL-# lines) as we selected these for exhibiting ambiguous (N’s) or anomalous (5′-CAT *→* 5′-CAG) DGRP assemblies. To better understand the distribution of *opa* variants in DGRP lines with unambiguous assemblies, we therefore decided to obtain and genotpe two additional sets of 12 lines chosen by one non-random strategy and one random strategy.

A recent report on radiation resistant RAL lines (Vaisnav *et al.* 2014) listed four of the Notch Dirty Dozen lines, which we had found to carry non-*opa31* variants, as exhibiting different degrees of radiation resistance (Table 1). We speculated that extremely variant *opa* lengths may lead to proteostatic stress. Aberrant pQ-related aggregation might induce the activity of stress response pathways and prime such lines to resist lethal doses of radiation (more on the phenotypes of *opa* variants later). In order to test this hypothesis, and obtain additional and more random sampling of the DGRP distribution, we genotyped 12 of the top radiation resistant lines (66/161) and 12 randomly-chosen radiation-sensitive lines (RAL 820–911), which constitute the bulk (∼60%) of the DGRP stocks (95/161). Altogether, we find that almost 62% of the radiation-resistant lines have non-*opa31 N* genotypes versus 29% of the radiation-sensitive lines (Table 1). Furthermore, the average absolute value of deviation from the wild-type length of 31 amino acids is 1.9 amino acids for the radiation-resistant lines, whereas it is less than 0.6 amino acids for the radiation-sensitive lines (Table 1). Thus, non-wild-type *N opa* length variants are strongly correlated to radiation-resistance. We speculate as to whether this may involve stress in the endoplasmic reticulum (ER), where the Notch membrane receptor is translated and extensively processed (Fortini 2009; Yamamoto *et al.* 2010). Up-regulation of the unfolded protein response (UPR), which is induced by misfolded proteins in the ER, is known to confer treatment-resistance to some cancers (Nagelkerke *et al.* 2014).

The strange RAL-646 stock, which is homozygous for *opa23 N* on the X chromosome, is one of only three RAL lines that are also homozygous for the Kodani inversion *In(3R)K*, located in the right arm of chromosome 3 (Huang *et al.* 2014). Ten additional lines are heterozygous for *In(3R)K*, including RAL-440, which we genotyped as having wild-type (*opa31*) repeats. The Kodani inversion is a rare but cosmopolitan inversion that originated about 20–100 thousand years ago, although it was recently discovered that it is almost entirely fixed in the African samples from the Oku range in Cameroon (Corbett-Detig and Hartl 2012). Because of the interesting *opa23*-related phenotypes described later, we also genotyped the *opa* repeats of the two other lines that are homozygous for the Kodani 3R inversion, RAL-100 and RAL-105. We find that RAL-100 is homozygous for *opa32*, while RAL-105 is homozygous for *opa33b*. Of these genotypes, *opa33b* figures prominently in our discoveries concerning *w*^*a*^-linked X balancer chromosomes (described in a separate section below).

In Figure 2, we show all of the 40 RAL lines we genotyped as a single distribution. This distribution shows the frequencies of 10 different Notch *opa* isoforms sampled in the Notch Dirty Dozen (ambiguous DGRP1.0 sequence information), 12 additional radiation-resistant lines, 12 additional radiation-sensitive lines, and two additional lines homozygous for *In(3R)K* (individual lines per group are listed in Table 1). This distribution has three remarkable features (Fig. 2). First, nearly 49% of examined RAL lines (19.5/40) do not have wild-type *opa31* repeat haplotypes and are thus not being accurately genotyped by current high-throughput methods. Second, it shows that this allelic distribution is extremely asymmetric. There is a precipitous drop in frequency for alleles longer than *opa32* but nonetheless there is a long tail of many rare *opa* expansion alleles in the *opa33–opa37* range. Third, there is a deficit of alleles shorter than the wild-type *opa31* genotype except the *opa23* variant, which is also missing the intervening histidine. Thus, the *opa24–opa30* range, which includes variants up to 7 residues shorter than wild type, is a veritable “allelic barrens” for *D. melanogaster*. This allelic barrens is in stark contrast to an “allelic jungle” of ten distinct rare *opa* variants spanning the range up to 7 residues longer than wild type *opa31*.

#### Confirmation of the opo23 genotype for RAL-646

The DGRP “Line-646” assembly for RAL-646 was reported as a single H → Q variant, which is what originally garnered our attention. Instead, we found this line to be homozygous for *opa23*, which is 24 bp shorter than the reference *opa31*. In retrospect, we find that the Line-646 DGRP assembly is a perfect example of the problematic nature of genotyping the *opa* sequences and of the repeat assembly problem in general. This problem is exacerbated by the need for flanking, non-repetitive “anchor” sequence to accurately start or end the repeat alignments in the correct position. However, the need for such anchor sequences in the read further reduces the remaining read length that can span into the repeats. For the *Notch opa* repeats, a single 75 nucleotide (nt) read cannot span the wild-type length of 93 bp even without anchor sequence. From what we have seen, 91% of *opa* length variants (10 of 11 observed variant length classes) have a greater number of repeats than wild-type. Nonetheless, even the shorter RAL-646 *opa* genotype must be assembled with anchored repeats. If a minimum of two flanking non-CAX codons are used for anchoring sequence, this leaves only 23 repeat CAX codons in a 75 nt read (6 + 69), and thus the read still does not bridge safely to anchor sequence on the other side of the repeats. We expand upon this example in greater detail below to explain how we ruled out other explanations for the Line-646 DGRP assemblies.

After determining a homozygous *opa23* genotype for RAL-646 (one of many lines we genotyped), we asked whether this line had changed since the time of the DGRP sequencing and assembly. For example, it could have been the case that an original change of the intervening His to Gln resulted in a toxic uninterrupted pQ tract. This non-synonymous mutated allele could have been replaced in the stock by a subsequent more advantageous deletion that shortened this tract. To address this question, we obtained all of the original DGRP sequencing reads and aligned them to the iso-1 reference alignment (Supporting File S1). Many of the reads in this alignment had mismatches to each other and to the reference sequence, and a few only had one or two that were plausibly polymorphisms specific to this stock. After removing the reads with high levels of mismatch, we are left with an alignment that suggests the two synonymous changes and the indicated non-synonymous change seen in the Line-646 DGRP assemblies. It would have been computationally challenging to consider all alignments in the space encompassing indels up to 24 bp. Nonetheless, when we re-assemble the original DGRP reads by alignment to our long Sanger sequence, we increase the number of original DGRP reads with 100% alignment (Supporting File S1). Thus, the original RAL-646 stock was and continues to be homozygous for the unique *opa23* allele and not the reported substitution variant encoding an H → Q *opa31*. This typical DGRP assembly artifact, which is inherent to short read assemblies, is probably exacerbated by the intractability of considering a larger space of possible indels, an over-reliance on alignment to reference genome, and the reference iso-1 *opa31*2* genotype, which does not conform exactly to the common sequence of CAG and CAA codons. In summary, we find that the important *N opa* repeats are best sequenced and assembled using sequence reads > 140 bp long (3 bp × 40 triplet repeats + 10 bp × 2 anchor sequences).

### 3. Classic N mutant alleles do not harbor opa length diversity, unlike X balancers

The slightly expanded *opa32* version of *N* was previously reported for the classic alleles *N*^*Co*^ and *N*^60*g*11^ but this extra glutamine is thought to be phenotypically silent (Kidd *et al.* 1986; Lyman and Young 1993). We set out to survey the *opa* repeat genotypes for additional *N* alleles with unknown *opa* repeat configurations on the supposition that some classic mutants may have acquired suppressors via changes in the *opa* repeats after decades of maintenance at stock centers. Remarkably, given the many new *opa* variants we had found by surveying DGRP stocks, our survey of *N*^1^, *N*^*ts*^1, *N*^*ts*2^, *N*^*nd*0^, *N*^*TAV17V*^, and *N*^*Ax-ts*1^ did not turn up a single length variant different from *opa31* (Figs. 2 and 3). This striking result suggests to us that perhaps changes in the *opa* repeat number by contraction or expansion is generally deleterious in combination with existing lesions at this locus.

In the process of genotyping the *N*^1^ and *N*^*TA17V*^ stocks, which are kept over the multiply inverted *FM7c* balancer chromosome, we discovered that this and most other X balancers had the rare *opa33b* allele, and two related X balancers had *opa32*. We found no X balancers carrying the major *opa31* alleles. It is important to note that these balancers all involve the *In(1)sc*^8^ inversion, which effectively reduces the dosage of *AS-C* expression in proneural clusters and in the sensory organ precursor cell itself [reviewed in (García-Bellido and De Celis 2009)]. We subsequently genotyped the *In(1)sc*^8^ stock and confirmed that it contained the *opa33b* allele. This inversion was produced by Sidoroff in 1929 by X-ray mutagenesis of a fly stock carrying the *apricot* allele of *w*, which is linked to *N* by 1 cM (Sidorov 1931). Thus, we find that all *w*^*a*^-carrying balancers that we genotyped (*FM1*, *FM7a*, *FM7c*, and *Basc*) carry the *opa33b* allele, while those balancers that no longer carry *w*^*a*^, *FM4* and *FM6*, are linked to *opa32*. Later, we present experimental evidence consistent with suppression of the *sc*^8^ inversion phenotype by the expanded *opa33b* allele of *N* carried within the inversion. These results also show that the *opa* variants can be stably maintained for hundreds of generations. We suspect that this involves selection as we have found that rare variant *opa* length alleles from the RAL stocks produce specific developmental phenotypes.

### 4. Short and long N opa variants cause developmental defects

We noted that the RAL-646 stock, which is homozygous for the short *opa23* rare variant allele, was much more difficult to expand than many other RAL stocks homozygous for the wild-type *opa31* allele. We attribute this in part to a high rate (27%) of embryonic lethality in the RAL-646 stock (Table 2 and Figure 4A). In comparison, seven other stocks that were either homozygous or heterozygous for the two most common alleles, the wild-type *opa31* and the related *opa32*, had an average of 13.1% embryonic lethality (*i.e.*, < 1/2 RAL-646 lethality) when grown at room temerature (22–23*°*) or 25*°*. Interestingly, the RAL-237 stock, which carries the long *opa35* rare variant allele, had the lowest embryonic lethality at 3.7%.

**Table 2.** RAL inbred lines with the shortest and longest opa variants have the highest and lowest lethality, respectively

| RAL line<sup>a</sup> | Genotype | Embryonic lethality | Assay temp. | Fecundity at 25°<sup>b</sup> |
| --- | --- | --- | --- | --- |
| 646 | *opa23* | 27.1% | 25° | 1,436 eggs (63.1%) |
| 822 | *opa31* | 20.9% | 25° | 1,438 eggs (63.2%) |
| 439 | *opa31* | 20.3% | 25° | 1,699 eggs (74.6%) |
| 142 | *opa31/opa32* | 15.9% | 22–23° (R.T.) | n. d.<sup>c</sup> |
| 21 | *opa31* | 15.4% | 25° | 1,196 eggs (52.5%) |
| 440 | *opa31* | 9.1% | 25° | 742 eggs (32.6%) |
| 338 | *opa32* | 5.2% | 22–23° (R.T.) | n. d. |
| 208 | *opa31* | 5.2% | 22–23° (R.T.) | n. d. |
| 237 | *opa35* | 3.7% | 25° | 2,276 eggs (100%) |
| AVG. |  | 13.7% |  | 1,465 eggs (64.3%) |

<sup>a</sup> Lines are ordered from the most to least embryonic lethality
<sup>b</sup> Number of eggs laid in 28 hour period beginning 2 h before dusk. Average fecundity was 1,465 eggs (64.3% of maximum 2,276).
<sup>c</sup> not determined

**Figure 4.**
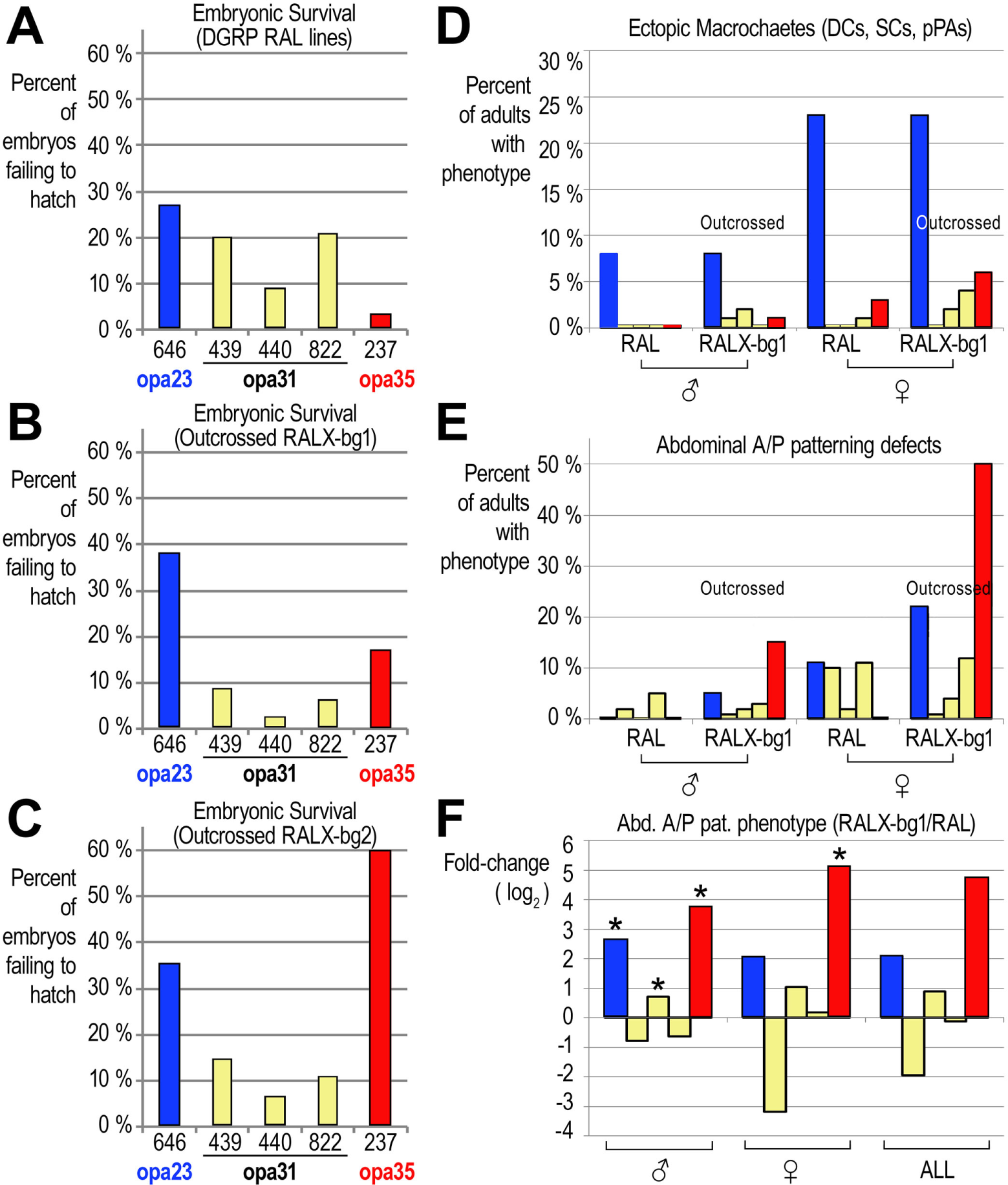
RAL-X chromosomes carrying extreme *opa* length variants display common phenotypic effects upon out-crossing. **(A)** Embryonic lethality is high in the isogenic RAL-646 line, carrying *opa23*, while it is lowest in the isogenic RAL-237 line carrying *opa35a1*. Three different isogenic RAL lines carrying wild-type *opa31* sequences have intermediate ranges of embryonic lethality. **(B)** In outcrossed RAL X chromosome lines (RALX-bg1), embryonic lethality is increased in the extreme *opa* variants, while it is decreased in all of the wild-type *opa31* lines. To produce the RALX-bg1 lines, we used *FM7c*/Y males from a lab stock of *N*^1^/*FM7c* (background one, or bg1) to outcross the RAL X chromosome five generations. The RAL X chromosome was then re-homozygosed after two more generations of sib-crosses to lose the balancer X chromosome. **(C)** This is similar to panel B, except RALX-bg2 lines were established using males from a lab stock of FM7a (background two, or bg2). In this background, both of the outcrossed RAL X chromosomes from the parent lines with extreme *opa* length variants RAL-646 (*opa23*) and RAL-237 (*opa35a*) have severe embryonic lethality. RALX237-bg2 is particularly more pronounced in this defect relative to RALX237-bg1. **(D–F)** Similar results with other phenotypes involving the same six RAL X chromosomes from inbred and outbred DGRP lines are shown (left to right: RAL-646, RAL-439, RAL-440, RAL-822, RAL-237). **(D)** A similar intensification of phenotype in the derivative RALX-bg1 lines carrying *opa* length variants is seen for ectopic macrochaete (doubled bristles) on the adult thorax (DCs, dorsocentral macrochaetes; SCs, scutellar macrochaetes; pPAs, posterior post-alars). This defect is most pronounced for the RALX646-bg1 line but some intensification is also seen for the RALX237-bg1 line. Females from both lines are more affected than males, suggesting a sensitivity to the specific X dosage. (Male X-linked genes are dosage-compensated, but not always to the exact same levels as the female gene, with some variation for different X-linked genes.) **(E)** A similar intensification of an abdominal anterior/posterior (A/P) patterning defect (see supplementary Fig. S) is seen in the derivative RALX-bg1 lines carrying *opa* length variants. This defect is also more pronounced in females. **(F)** The same data as in E, transformed to show fold-difference of the outcrossed RALX-bg1 lines over the parent RAL lines. The columns with asterisks represent ratios in which a pseudo-count of 1.0 is used for the phenotypic values of the RAL parent lines to avoid division by zero.

We also measured fecundity at 25*°* (# of eggs laid in a 28 h period, dusk-to-dusk). This shows that the RAL-646 stock has a nearly average fecundity, demonstrating that this stock’s failure to thrive stems mainly from its increased embryonic lethality, possibly caused by its *opa23 N* sequence. Of note, the inbred RAL-237 line again outperformed all other stocks in this regard (Table 2).

### 5. Effects of outcrossing RAL N opa variants

To investigate whether the RAL-646 and RAL-237 lines were exhibiting outlier phenotypes in the embryonic lethality assays due to their variant X-linked *opa* repeat sequences, we assayed this phenotype and others after outcrossing diverse RAL lines to a common background. For RAL-646 (*opa23*), RAL-237 (*opa35*) and three *opa31* lines (RAL-439, RAL-440, and RAL-822), we crossed virgin females with balancer X males from a “background” stock of *FM7c/N*^1^ (“bg1”) or *FM7a* (“bg2”). The *FM7a* background (bg2) was chosen for comparison to *FM7c*, which possibly could have acquired *N*^1^ suppressors. For four additional generations we took virgin females and crossed them back to the balancer males, followed by re-isogenizing the RAL X chromosomes to produce a series of RALX-bg1 and RALX-bg2 5x outcrossed lines. For comparison, we assayed embryonic lethality for each common background series in parallel with the RAL parent stocks.

We find that embryonic lethality intensifies significantly for *both* the *opa23 and* the *opa35* outcrossed lines, while it becomes attenuated in *all* six of the *opa31* lines (Fig. 4B, C). In fact, while the *opa35* RAL-237 stock had the lowest embryonic lethality in all experiments, the RALX237-bg2 had the highest embryonic lethality at 60%. These results suggest that the predominant major allele *opa31* does best in outcrossed lines, while the *opa35* can do best in an isogenized bacground with compatible (suppressor) modifiers. In contrast, these results suggested that the *opa23* allele is likely deleterious in most genomic backgrounds. Below, we show that these effects and other classic *Notch* phenotypes intensify even more when we recombine away the *N*^*opa23*^ allele from the RAL-646 X chromosome.

We also measured the frequency of duplicated macrochaetes on the notum of adult flies, a classic *N* phenotype, because the RAL-646 (*opa23*) parent line had increased numbers of ectopic macrochaetes in both males (8%) and females (23%) (compare Fig. 5A–C). We scored the frequency of ectopic dorsocentral (DCs), scutellar (SC) and posterior post-alar (pPA) macrochaete-type bristles (Fig. 4D). We find that these numbers do not change much for the outcrossed RALX646-bg1 line. This phenotype did intensify in the RALX237-bg1 line although not nearly to the same levels as the RAL-646 or RALX646-bg1 lines (Fig. 4D). Nonetheless, the two outcrossed lines carrying *opa23* or *opa35* have the highest levels of ectopic bristles in females, >3x higher than outcrossed males. This suggests that this phenotype is sensitive to precise X dosage levels (XX vs. male dosage compensated X).

**Figure 5.**
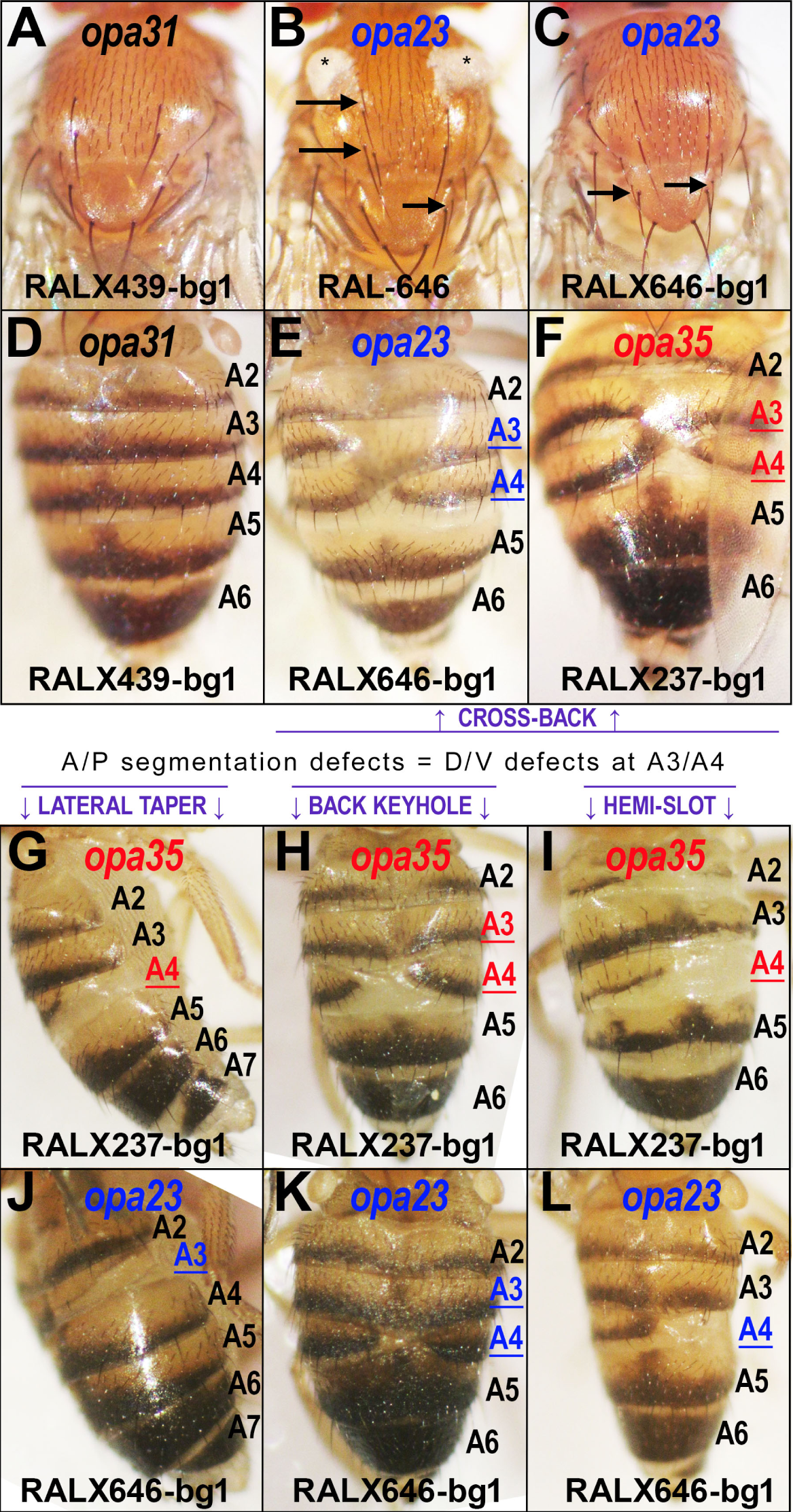
Both short and long *opa* length variants produce common phenotypic effects in adult flies. **(A–C)** Macrochaete patterning in females of various parent and outcrossed RAL lines. **(A)** The outcrossed RALX439-bg1, a typical *opa31* line, is shown as a wild-type representative. **(B, C)** Both the RAL-646 **B** and outcrossed RALX646-bg1 **C** flies have a high frequency of forming ectopic macrochaete bristles (arrows). Symmetrical cuticle fluff also appears in the parent RAL-646 (asterisks in **B**) but is not present when outcrossed. **(D–F)** Abdominal anterior/posterior (A/P) patterning in adult females of various parent and outcrossed RAL lines. **(D)** The outcrossed RALX439-bg1 is shown as a typical wild-type *opa31* control. **(E–L)** Both the outcrossed RAL X chromosomes carrying either a short *opa23* (**E**, blue) or long *opa35* (**F** red) allele have a high frequency of abdominal A/P patterning defects, including the “crossback” phenotype shown (**E, F**) in which pigmented stripes from two segments appear to cross-over each other. In addition to the cross-back defect, other A/P segmentation defects include the lateral taper (**G, J**), the back keyhole (**H, K**), and the hemi-slot (**I, L**). All of these defects predominantly involve the A3 and A4 segments (highlighted and underlined) of the RALX-bg1 lines carrying the shorter *opa23* (E, J–L) or longer *opa35* (F–I) alleles.

We saw a pattern of phenotypic intensification after outcrossing for one other phenotype pertaining to dorsal–ventral (D–V) patterning along the anterior–posterior (A–P) abdominal segments (Fig. 4E–F, and Fig. 5D–L). This phenotype can be seen in many different RAL lines at low frequencies (1–10%) but is typically more frequent in females than males. After outcrossing, this phenotype significantly increases in both male and female *opa35* RALX237-bg1 flies with a higher penetrance in female flies (Fig. 4E–F). These D–V/A–P phenotypes can be divided into at least four related types, which we refer to as the “cross-back” (Fig. 5E, F), “lateral taper” (Fig. 5G, J), “back keyhole” (Fig. 5H, K), and “hemi-slot” (Fig. 5I, L) phenotypes. Examples of all of these D–V/A–P defects can be found in both the *opa23* RALX646-bg1 and *opa35* RALX237-bg1 lines. These phenotypes predominantly affect D–V patterning axis at the A3 and A4 abdominal segments. A comparison of fold-change (outcrossed/inbred) shows that these D–V/A–P defects typically decrease for the lines carrying *opa31* while they increase manifold for the lines carrying length variants *opa23* and *opa35* (Fig. 4F).

### 6. Aberrant opa lengths affect rho expression along the embryonic A–P axis

During embryonic D–V patterning, Notch signaling is known to act upstream of EGFR signaling via the *rhomboid* locus (*rho*) (Markstein *et al.* 2004; Crocker *et al.* 2010). We thus performed in situ hybridizations with an anti-sense RNA probe to determine *rho* expression patterns in the early embryo. We find that many RALX646-bg1 embryos could be found in which the A–P modulation of the lateral stripe of expression does not resolve as cleanly as it does in control RALX440-bg1 embryos (Fig. 6). In fact, if we look at later embryonic stages through long-germ band extension, we can find many RALX646-bg1 embryos with either missing or disrupted *rho* expression along the posterior edges of the ventral midline. Thus, we suspect that the rare contracted or expanded pQ versions of Notch critically impact embryonic N signaling through the *rho* locus via one or more of its many conserved enhancer modules (we are currently pursuing partially-redundant enhancers acting later than the *rho* NEE).

**Figure 6.**
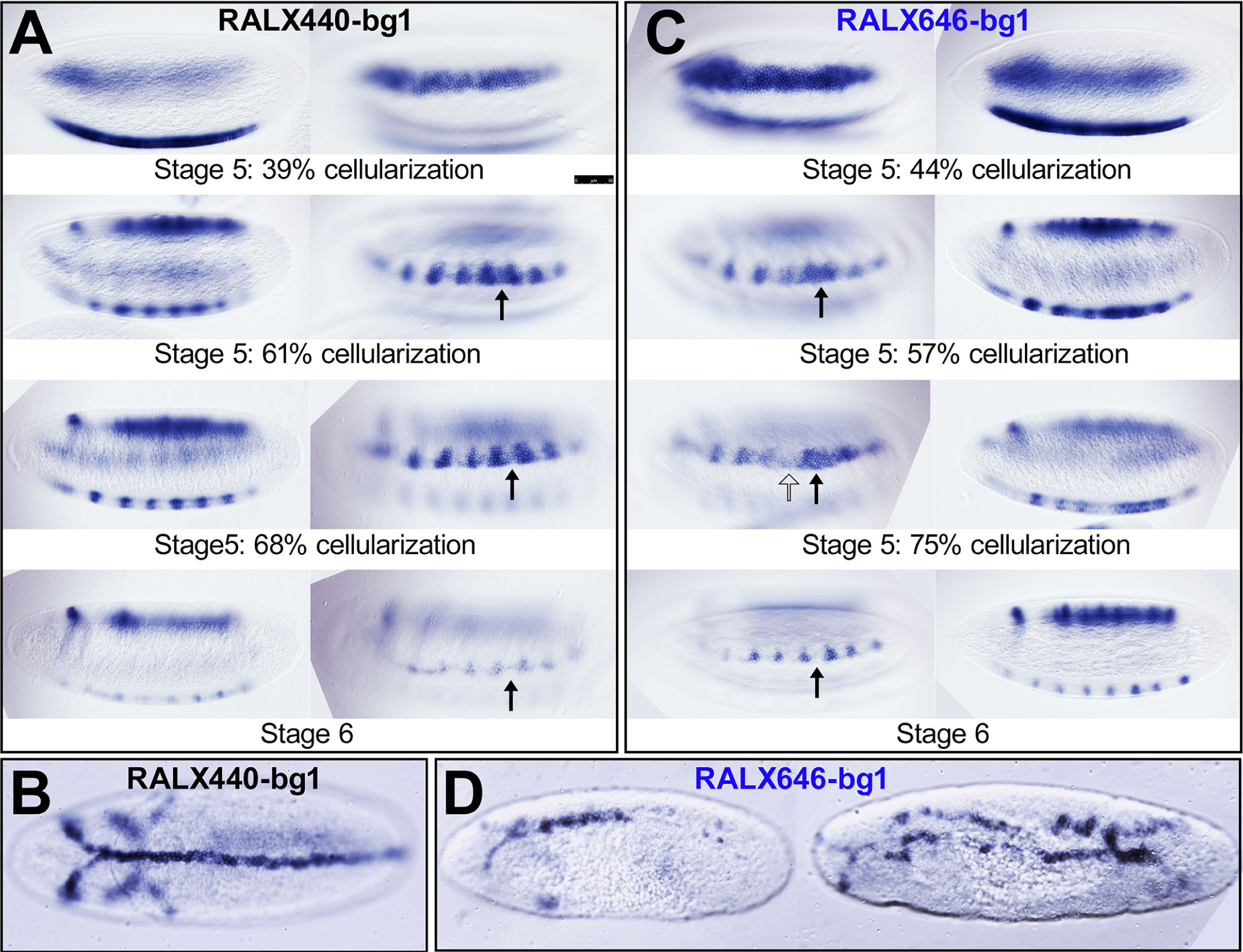
The anterior–posterior (A–P) modulation of the dorsal/ventral (D/V)-specified lateral stripe of expression for *rho* is affected by *opa* variants. **(A, C)** Stage 5–6 embryos stained for *rho*. In each row of each panel is a pair of images for the same embryo. The outer (side) embryo is an optical cross-section used to stage the degree of cellularization. The inner embryo is a surface view showing the lateral stripe of *rho* expression. **(A, B)** Expression of *rhomboid* (*rho*) mRNA transcripts via in situ hybridization with a digU-labeled anti-sense RNA probe against *rho* in Stage 5–6 (**A**) and Stage 10 long-germ band extended (**B**) embryos from the outcrossed RALX440-bg1 line carrying an *opa31* wild-type *N* locus. **(C, D)** Expression of *rho* mRNA transcripts in Stage 5–6 (**C**) and Stage 11 long-germ band retracting (**D**) embryos from the outcrossed RALX646-bg1 line carrying a deleterious *opa23 N* locus. Relative to wild-type *rho* expression, these embryos exhibit a higher probability of failing to refine the NEE-driven lateral stripe of expression in cellularizing Stage 5 embryos, into a series of well-separated triangular patches of expression (compare arrows in **A** versus **C**). As the early *rho* NEE-driven reporter activity fails to also drive the later triangle enhancer pattern, we postulate the existence of additional, late-acting *rho* enhancers that integrate both D/V and A/P signaling inputs.

N-Delta is important for A/P segmentation in vertebrates (Morales *et al.* 2002) and many arthropod groups such as spiders (Stollewerk *et al.* 2003), centipedes (Chipman and Akam 2008), and crustaceans (Eriksson *et al.* 2013). Based upon our findings in *Drosophila*, Notch may play a mixed role in A–P (not A/P) posterior morphogenetic patterning of holometabolous insects as it does in hemimetabolous insects such as the cricket (Mito *et al.* 2011). Interestingly, our results suggest that such roles may be limited to specific pQ-sensitive N signaling pathways.

To understand the nature of the extreme augmentation of the D–V/A–P segmentation defect in the outcrossed RALX237-bg1 line, which carries an expanded *opa35* variant, we sequenced a small selection of genomic regions encoding other potentially variable pQ tracts that might interact with NICD in the early embryo. Specifically, we Sanger re-sequenced a few prominent pQ-encoding repeats at *mastermind* (*mam*), *dorsal* (*dl*), and *twist* (*twi*) from several RAL lines.

Here, we report finding an expanded pQ-encoding tract at *dorsal* that may be relevant. We find that 72.7% of chromosome 2L haplotypes from RAL lines that are homozygous for *opa* length variants *≥* 32 encode a variant Dorsal transcription factor protein featuring the pQ tract Q_3_HQ_16_ and more extended versions (Table 3). In contrast, 0.0% of 2L haploytpes from RAL lines homozygous for *opa* length variants *≤* 31 encode this type of pQ tract at *dorsal*. Instead, they encode the major allele pQ tract of Q_3_HQ_14_, which is shorter in length by two glutamines. Interestingly, RAL-405, which is homozygous for *N opa31* on the X chromosome, is heterozygous for Q_3_HQ_14_ and the interesting allele Q_3_HQ_7_HQ_8_. The latter RAL-405 dorsal haplotype appears to be a Q_3_HQ_16_ that has undergone a non-synonmyous change of a His → Gln in the center of the long pQ tract. This change is consistent with compensatory selection for reversal variants of the longer expanded *dorsal* allele in an *opa31* background. We point out that Dorsal is only one of many pQ-rich factors that cotarget a small set of enhancers also targeted by Su(H) and NICD. Thus, we remain open to the possibility of finding additional, potentially-relevant, pQ modifiers. We address this and related findings in the discussion.

**Table 3.** Correlated pQ-encoding genotypes at Notch (chr. X) and dorsal (chr. 2L)

| *N opa* repeat # | Q<sub>3</sub>HQ<sub>14</sub> dorsal | Q<sub>3</sub>HQ<sub>16</sub> dorsal<sup>a</sup> | RAL lines genotyped by Sanger sequencing |
| --- | --- | --- | --- |
| ≤ 31 | 94.4% (8.5/9 lines) | 0.0% (0/9 lines) | 21, 91, 208, 350, 405<sup>b</sup>, 439, 440, 646, 822 |
| ≥ 32 | 22.7% (2.5/11 lines) | 72.7% (8.0/11 lines) | 57<sup>**</sup>, 69<sup>**c</sup>, 88<sup>*d</sup>, 237<sup>**</sup>, 338<sup>**</sup>, 370<sup>**</sup>, 555, 776, 796<sup>**</sup>, 801<sup>**</sup>, 802<sup>*e</sup> |

<sup>a</sup> \* Lines with asterisks have this pQ haplotype
<sup>b</sup> This line is heterozygous for Q<sub>3</sub>HQ<sub>14</sub> and Q<sub>3</sub>HQ<sub>7</sub>HQ<sub>8</sub>.
<sup>c</sup> This line is actually heterozygous for Q<sub>3</sub>HQ<sub>21</sub> and Q<sub>3</sub>HQ<sub>20</sub>.
<sup>d</sup> This line is heterozygous for Q<sub>3</sub>HQ<sub>16</sub> and Q<sub>3</sub>HQ<sub>15</sub>.
<sup>e</sup> This line is heterozygous for Q<sub>3</sub>HQ<sub>16</sub> and Q<sub>3</sub>HQ<sub>14</sub>.

### 7. Variant opa lengths lead to macrochaete patterning defects

The developmental induction of proneural clusters for specific macrochaetes involves distinct pathways targeting a number of independent “pre-patterning” enhancers (PPEs) at the *ac sc* complex (*AS-C*) (Martínez and Modolell 1991; Skeath *et al.* 1992; Gómez-Skarmeta *et al.* 1995; Modolell and Campuzano 1998). These sensory bristle-specific regulatory modules and the *AS-C* gene copy number are also dynamically evolving (Galant *et al.* 1998; Negre and Simpson 2009). Importantly, mutations in genes for N-Delta signaling components can affect specific bristles. This may be because proneural cells transition from the effects of any number of diverse pre-patterning signals to the lateral inhibition regulatory network featuring N-Delta signaling. As the 26 symmetric macrochaete sensory organs on the adult notum are also monomorphic traits, these circuits must also function in the presence and absence of male dosage compensation of X-linked genes, including the *AS-C* and *N* genes. Thus, N-Delta signaling must become highly canalized in order to build the stereotypical macrochaete-type sensory organ at several positions receiving distinct signaling cues and operating at slightly different dosage levels. Some preliminary work has already been done to model parameter space across different mechanistic contexts (*e.g.*, lateral inhibition, boundary formation, asymmetric cell fate specification) (Matsuno *et al.* 2003; Barad *et al.* 2010, 2011; Sprinzak *et al.* 2011; Shaya and Sprinzak 2011).

To understand whether specific macrochaete bristles were being affected in adult male and female flies, we scored the relative presence and absence of each of the 13 macrochaete bristles of the adult hemi-notum in our RAL parent lines and 5x outcrossed RALX-bg1 and RALX-bg2 lines (Fig. 7). These results show that it is primarily the posterior post-alar bristle (pPA), anterior and posterior dorsocentral bristles (aDC and pDC), and the anterior scutellar bristle (aSC) that are ectopically produced in lines carrying rare, variant *N opa* alleles. These results will form a backdrop to further interesting and specific effects on macrochaete specification caused by these variants in various contexts.

**Figure 7.**
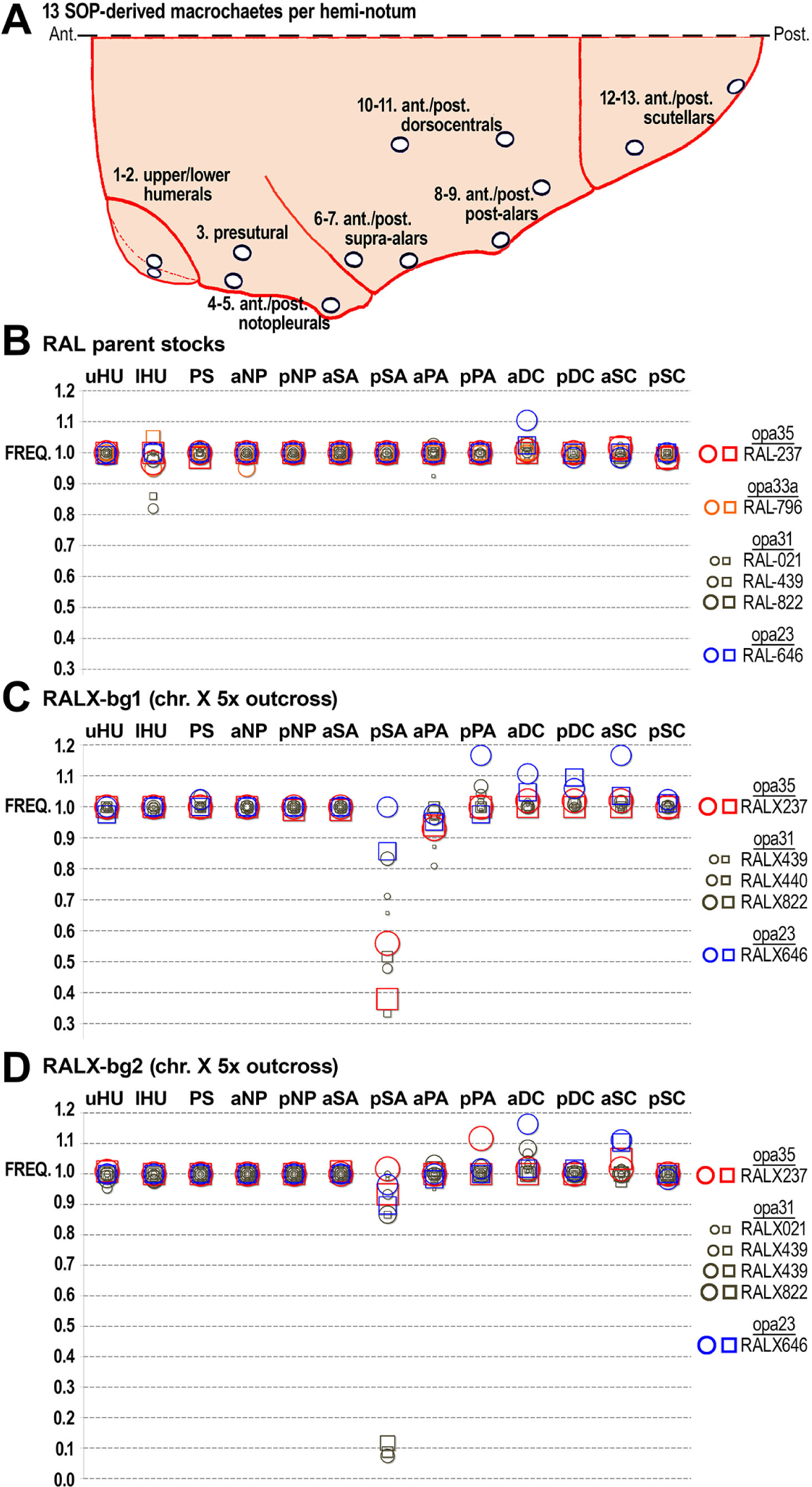
Macrochaete patterning defects in parent RAL versus outcrossed lines. **(A)** Cartoon of hemi-notum showing frequencies of all 13 macrochaetes of female (circles) and male (squares) flies from the indicated lines (colored shape outlines). In this and other figures, the macrochaete positions are abbreviated as follows: (1–2) upper and lower humerals (uHU, lHU); (3) presutural (PS); (4–5) anterior and posterior notopleurals (aNP, pNP); (6–7) anterior and posterior supra-alars (aSA, pSP); (8–9) anterior and posterior post-alaors (aPA, pPA); (10–11) anterior and posterior dorsocentrars (aDC, pDC); and (12–13) anterior and posterior scutellars (aSC, pSC). Anterior is to the left. Posterior is to the right. Top of image is the dorsal midline. **(B)** Frequency (FREQ) of macrochaete occurrence in parent RAL stocks. **(C)** Frequency of macrochaete occurrence in 5x outcrossed RALX stocks using *FM7c* males (bg1). Outcrossed lines are homozygous for the RAL X. The failure to form pSA is also seen in the *FM7* balancer lines (data not shown). **(D)** Frequency of macrochaete occurrence in 5x outcrossed RALX stocks using *FM7a* males (bg2). Outcrossed lines are homozygous for the RAL X. These data show that the RALX646 (blue *opa23*) and occasionally RALX237 (red *opa35*) outcrossed lines produce ectopic posterior post-alar, dorsocentral, and scutellar macrochaetes.

We also see a general failure to form the posterior supra-alar macrochaete in many of the outcrossed RALX-bg lines regardless of *N opa* status. Because we see a similar affect in *FM7* flies, and the 5x outcrossed lines underwent passage through these backgrounds before we re-isogenized the RAL X chromosomes, we attribute this affect to specific autosomal modifiers carried in the *FM7* genetic background.

### 8. N opa23 produces notched wings in males and females

In order to ascertain that the most extreme *N opa* length variants had phenotypes due to the *N opa* sequence and not to other loci on their X chromosomes, we recombined the interesting *N*^*opa23*^ allele into a different X chromosomal background (see Material and Methods). Our crossing scheme involved the parent RAL-646 line, and fly stocks of *w*^1118^ and *FM7* balancer stocks. The resulting white-eyed “wopa23” line carries a much smaller portion of the RAL-646 X chromosomal segment spanning from *N* to some cross-over point left of the *ABCF2* gene encoded by *CG9281* (Fig. 8A).

**Figure 8.**
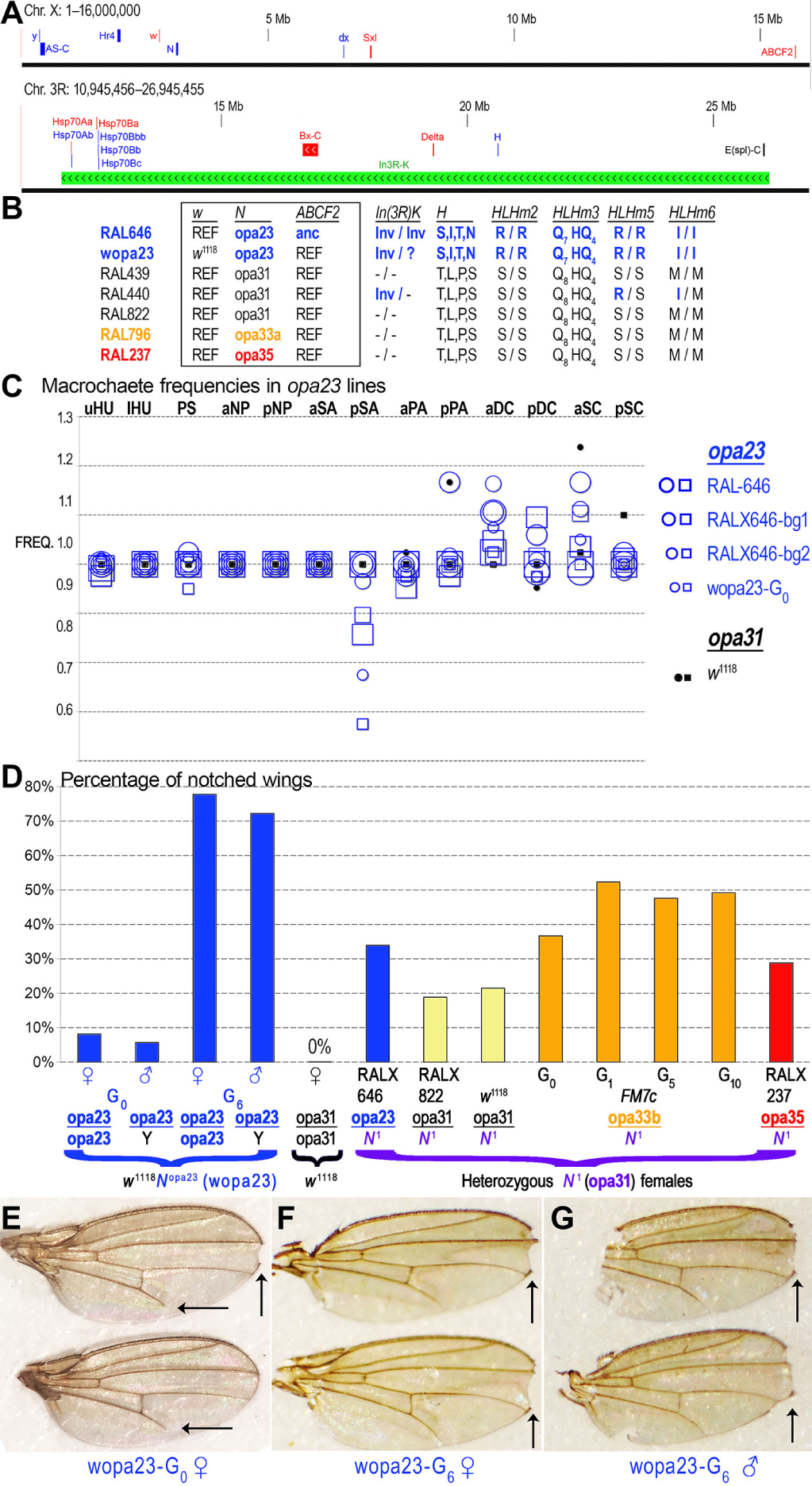
The deleterious *N opa23* variant produces notched wings after outcrossing. **(A)** Potential modifiers of *Notch opa* alleles. Shown are 16 Mb regions from chromosomes X and 3R. Genes indicated on the X chromosomal map include (left to right): *yellow* (*y*), the *achaete scute* complex (*AS-C*), *Hr4*, *white* (*w*), *Notch* (*N*), *Deltex* (*dx*), *Sex lethal* (*Sxl*), and *ABCF2* (*CG9281*). Genes indicated on the 3R chromosomal map include (left to right): *Hsp70* gene clusters, the *bithorax* complex (*Bx-C*), *Delta* (*Dl*), *Hairless* (*H*), the *Enhancer of split* complex (*E(spl)-C*), and the Kodani inversion *In(3R)K* (green bar), which is homozygous in RAL-646 and only two other RAL lines (RAL-100 and RAL-105). **(B)** Shown are various polymorphisms (substitutions, pQ variation, inversions) distinguishing RAL-646 from other RAL stocks or RAL-646-derivative lines. The boxed columns are X-linked, while the other columns correspond to loci inside the *In(1)sc*^8^ inversion. **(C)** Shown are the macrochaete frequencies for RAL-646, RALX646-bg, and wopa23 line in the same convention as previous figures. **(D)** The recombined RAL-646 X chromosome swaps out the RAL-X chromosome between *w* and *N* and to the left of *ABCF2* (*CG9281*). We refer to flies carrying this recombined X chromosome as the wopa23 line. The original “G_0_” generation (wopa23-G_0_) produces notched and/or nicked wings in males and females at a modest but not insignificant rate. However, following six generations of selection for notched wing flies, this rate increased significantly to 72% in males and 78% in females (wopa23-G_6_). In contrast, selection for notched wings in the *N*^1^/*FM7c* stock failed to increases the rate of notching past 53% after ten generations of selection. **(E)** Shown are a pair of wings from a *wopa23*-G_0_ female prior to selection for notched wings. Vertical arrow shows a classic wing notch. Horizontal arrows in each wing show an additional defect of incomplete L5 wing vein formation albeit at a lower frequency than notched wings. **(F,G)** Wings from female and male wopa23 flies after 6 generations of propagating only flies with notched wings. These flies have more deeply serrated notches than the unselected wopa23 stock. The incomplete L5 wing vein phenotype disappears in the wopa23-G_6_ flies, while the severity of wing notching increases.

As mentioned, the RAL-646 parent line is only one of three RAL lines that are homozygous for the rare but cosmopolitan *In(3R)K* Kodani inversion, which harbors the *Hsp70A* and *Hsp70B* clusters inside of one breakpoint, the *Bx-C* locus, *Delta*, *Hairless*, and the *E(spl)-C* just inside the second breakpoint (see Fig. 8A). Because this inversion harbors many Notch-signaling loci and a locus critical to posterior Hox patterning, we developed PCR assays to confirm and detect the *In(3R)K* inversion. This allowed us to confirm the *In(3R)K* homozygous genotypes for RAL-646, RAL-100, and RAL-105, and the heterozygous genotype for RAL-440 (Fig. 8B). Several protein coding polymorphisms within the Kodani inversion may be of interest to *Notch*-related phenotypes (Fig. 8B).

We scored macrochaete phenotypes for the wopa23 flies and compared them to our series of outcrossed *opa23* lines (Fig. 8C). This showed that male wopa23 flies have missing presutural macrochaetes about 5% of the time, which is a defect not seen in RAL-646, RALX646-bg1, RALX646-bg2, or *w*^1118^ flies of either sex. Elevated levels of ectopic aDC bristles are seen in the wopa23 flies, but these are comparable to the other *opa23* parent and outcrossed lines.

We find that our wopa23 male and female flies displayed an emergent classic *Notch* phenotype of notched and nicked wing tips (Fig. 8D, E). We also observed that a smaller number of flies had a wing vein patterning defect in which the L5 vein fails to reach the wing margin (horizontal arrows in Fig. 8E).

To understand the notched wing phenotype of the wopa23 flies better, we put various RALX-bg1 X chromosomes over an X chromosome carrying the dominant *N*^1^ allele, which produces notched wings at a certain frequency. We then scored the frequencies of notched and/or nicked wing tips in the F_1_ heterozygous non-balancer females. Our *N*^1^ cross results show that the *opa23* and *opa35* have the smallest suppression effect on notched wings in heterozygous F_1_ females. This suggests that both contractions and expansions of the wild-type *opa31* repeat length are types of lesions at the *N* locus.

We also performed an experiment on the *N*^1^/*FM7c* and wopa23 flies in which we selected for notched wings over several generations. While we increased the number of notched wings in *N*^1^ females modestly from 34% to 50% without being able to achieve further increases in notching, we substantially increased the rate of notched wings in both males and females from < 10% to > 70% in our wopa23 stock after only a few generations of propagating flies with notched wings (Fig. 8D). (The *N*^1^ allele is lethal in males, so we cannot score its effect on male wings.) Incidentally, these “Super-Notch” wopa23-G_6_ flies lose the L5 wing vein phenotype that we first saw with the wopa23-G_0_ generation, even while the severity of wing notching was increasing (compare Fig. 8E to 8F, G). We also re-genotyped the Kodani inversion in the Super-Notch wopa23-G_6_ stock and found that they still carried at least one copy of the *In(3R)K* inversion despite the three generations of outcrossing involved in the recombination screen. These results suggest the existence of certain NICD-targeted enhancers that play a critical role in the later stages of wing margin organizer activity and that are sensitive to NICD’s pQ tract lengths.

### 9. The sc[8] inversion carries a rare opa expanded Notch

We discovered that all X balancer chromosomes carrying *w*^*a*^ are linked to *N*^*opa33b*^ (Fig. 2). We trace this linkage back to Sidorov’s 1929 X-ray mutagenesis of *w*^*a*^ stock (Sidorov 1931), which produced the *sc*^8^ inversion allele *In(1)sc*^8^, which we genotyped and found to also carry *opa33b*. The *sc*^8^ inversion prevents diverse macrochaete pre-patterning enhancers and the sensory organ precursor enhancer from being able to drive both *achaete* (*ac*) and *scute* (*sc*) in the *ac*/*sc* complex (*AS-C*), which is indirectly downstream of Notch signaling via its repression by N-inducible *E(spl)-C*. Thus, the *opa33b* allele of *N* might represent an enhancer of the *sc*^8^ inversion that further de-canalizes robust N-Delta signaling. Alternatively, the *opa33b* allele might have been acquired and maintained as a suppressor of an otherwise decanalized N-Delta signaling caused by reduced dosages of Achaete and Scute factors.

To determine whether *opa33b* variant *N* suppresses or enhances the *sc*^8^ phenotype caused by a separation of diverse pre-patterning enhancers and SOP mother cell enhancer from *achaete* or *scute*, we identified the *sc* inversion allele that has the closest breakpoint to the *sc*^8^ breakpoint, *sc*^*V2*^. This *sc*^*V2*^ inversion allele has a breakpoint 6 kb upstream of the *sc*^8^ break but otherwise separates the same set of known pre-patterning and SOP enhancers present throughout the ∼105 kb long *AS-C* (Fig. 9). We genotyped this inversion allele and found it to encode wild-type *opa31* repeats at *N*. We therefore scored the presence and absence of macrochaetes in both inversion lines.

**Figure 9.**
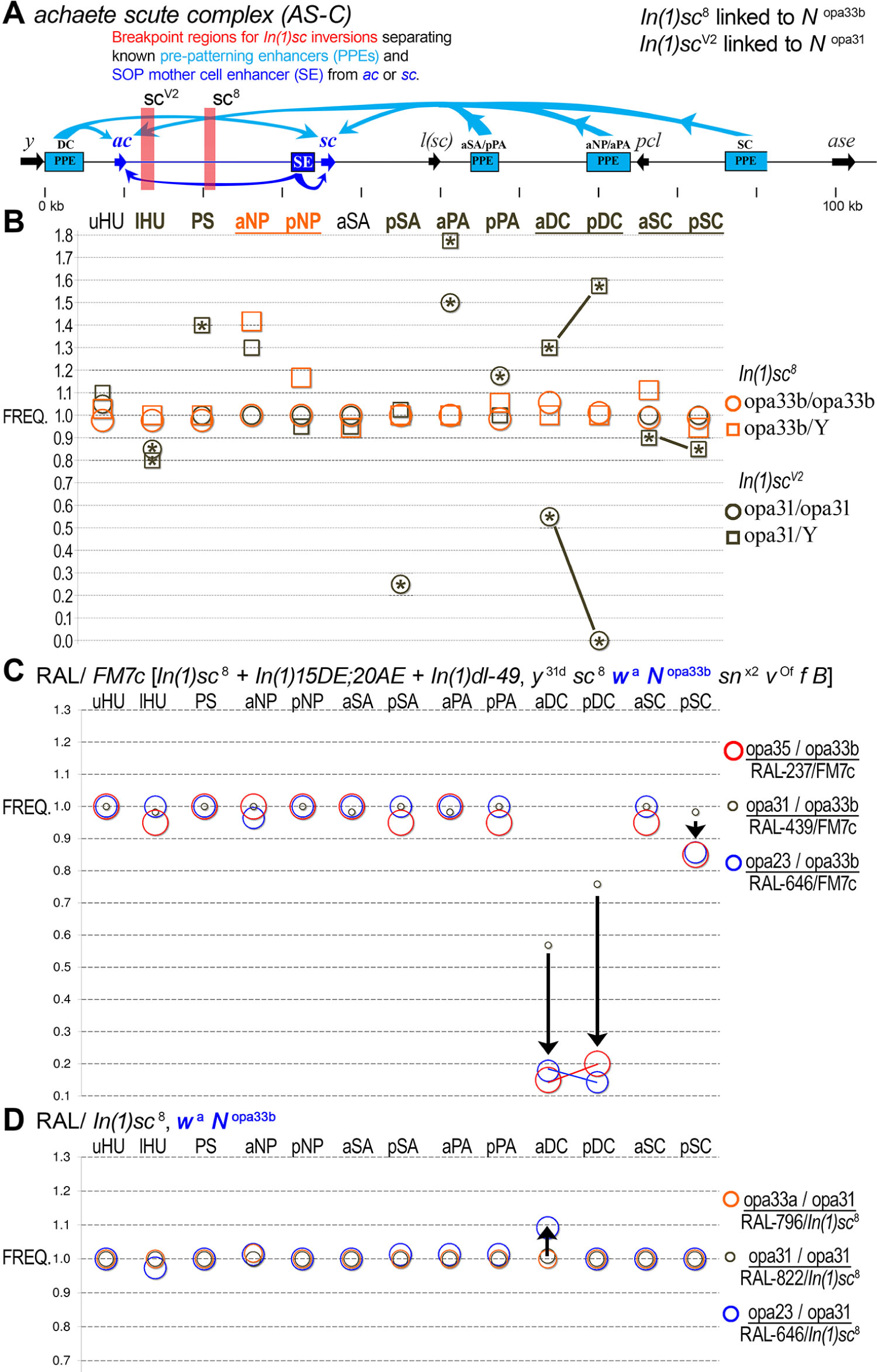
X balancers carrying the *white apricot* allele are linked to *N opa33b*. **(A)** Panel shows the *AS-C* locus and positions of the *sc*^*V2*^(linked to *opa31*) and *sc*^8^ (linked to *opa33b*) inversion breakpoints. Also shown are the diverse pre-patterning enhancers (PPE), which normally drive both *ac* and *sc* (blue arrows) in different proneural clusters of developing wing imaginal discs. In addition, the SOP mother cell enhancer (SE), which drives both *ac* and *sc*, is also shown. While each inversion is linked to a different *N opa* allele carried within the inversions, both inversions separate the same set of known enhancers. **(B)** A comparison of macrochaete patterning defects shows that *sc*^*V2*^has more deleterious effects than *sc*^8^, consistent with a suppressor role for *opa33b* expanded *Notch*. **(C)** A comparison of macrochaete defects for RAL stocks over *FM7c*, which carries the *sc*^8^ inversion allele. **(D)** RAL stocks over the *sc*^8^ inversion allele by itself, showing more modest effects.

We find that at several macrochaete positions, the *sc*^*V*2^line, which is linked to *N*^*opa*31^, has a more dramatic and penetrant phenotype compared to the *sc*^8^ inversion, which is linked to *N*^*opa*33*b*^ (Fig. 9). For example, *sc*^*V2*^ males have extremely high rates of ectopic DCs while *sc*^*V2*^ females have high rates of DC patterning defects. Empty sockets are not observed in females, which suggests that the defect is in under-specification of sensory organ precursors in females and over-specification in males. In stark contrast, *sc*^8^ males and females have almost normal DC bristle patterning in both males and females. This result is consistent with a role for the *opa33b* allele of *N* in partially suppressing *AS-C* insufficiency caused by the inversion’s localization of enhancers away from *ac* or *sc*.

To further test the hypothesis of *opa33b* functionality, we crossed our RALX-bg1 lines to *FM7c/N*^1^ flies and scored the heterozygous balancer females (RALX/*FM7c*). We find that the *opa23* and *opa35* heterozygotes are similarly severely deficient in the formation of DC and posterior SC macrochaetes relative to *opa31* heterozygotes (Fig. 9C). At several other bristle positions, the *opa35* (lHU, pSA, pPA, aSC) and *opa23* (aNP) variants are modestly affected relative to *opa31* heterozygotes. We think this result is significant and meaningful. However, similar experiments with RALX/*In(1)sc*^8^ heterozygotes gave us very different results, although *opa23* heterozygotes continue to exhibit outlier phenotypes, albeit more modestly than before. This suggests that the *In(1)sc*^8^ stock and the *FM7c* stock, which includes the *In(1)sc*^8^ inversion in addition to two other X inversions and several mutational markers, possess divergent sets of modifiers. Thus, the *N*^1^/*FM7c* balanced stock may have acquired and/or maintained suppressor modifiers of the *N*^1^ phenotype.

### 10. Diminished levels of opa35 Notch protein

To make sense of the phenotypic effects caused by pQ-variant Notch protein, we used a monoclonal antibody specific to the Notch intracellular domain (NICD) to compare levels of Notch full-length (N-FL) and cleaved NICD during the early embryonic stages corresponding to the mis-regulation of *rhomboid*. We first performed Western immunoblot for different time points in embryogenesis, which shows that the 0–2 hour embryos (0–2 h after egg deposition) have only full-length Notch, while later time points (2–4 and 4–6 h AED) have increasing amount of cleaved NICD (Fig. 10A). When we perform similar experiments comparing the *opa31* RALX822-bg1 line to the *opa35* RALX237-bg1, we find that there are greatly reduced levels of full-length N and NICD at 4–6 hour in the RALX237-bg1 embryos (Fig. 10B). One possible explanation for these observations is increased degradation of the Notch protein in RALX237-bg1 embryos, which in turn could lead to at least two of the phenotypic effects observed in this line and others.

**Figure 10.**
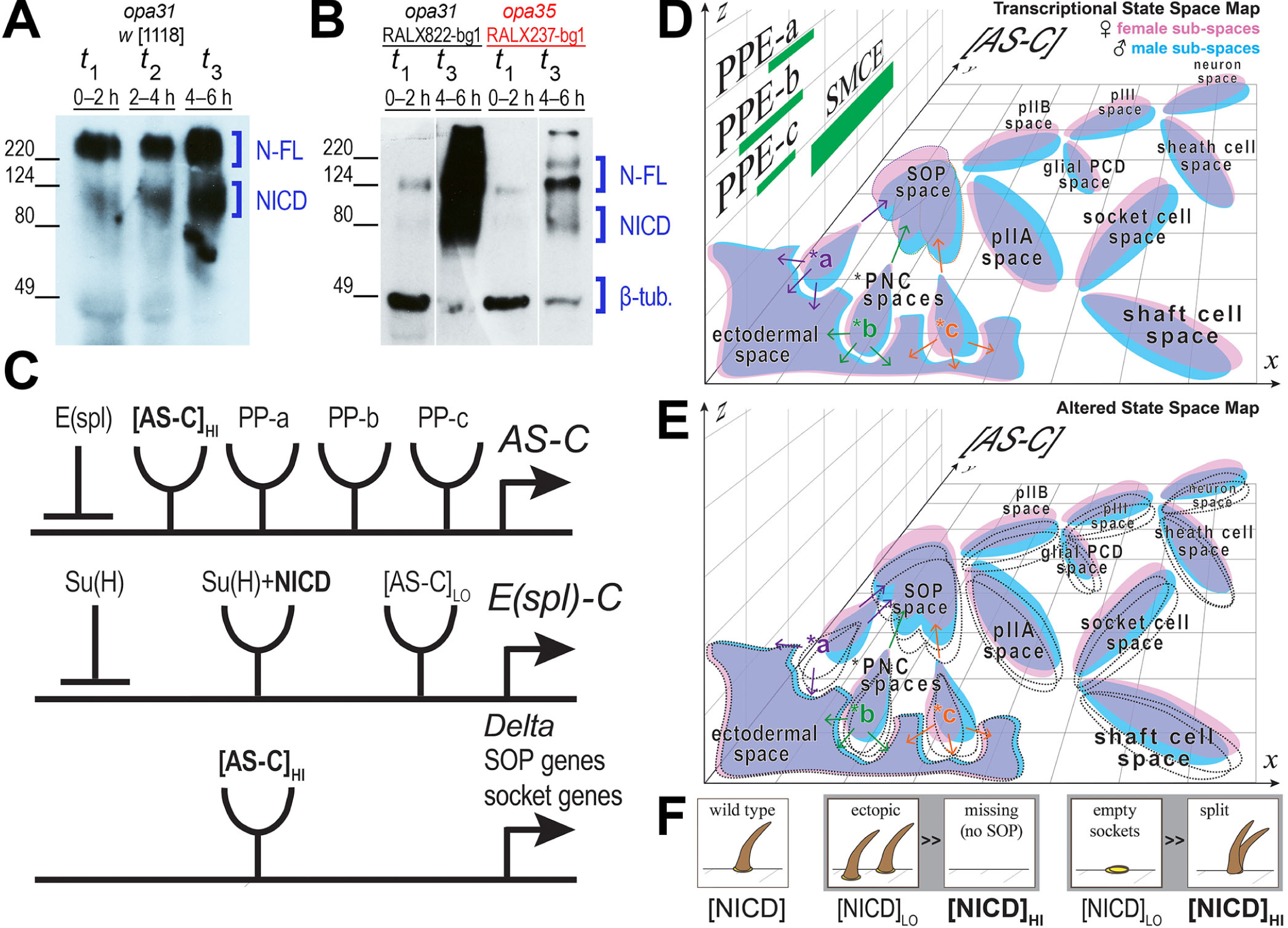
Extreme *opa* length variants appear to have lower levels of N protein, consistent with phenotypic outcomes caused by increased levels of Ac and Sc. **(A)** Western blot with an NICD-specific monoclonal antibody (DSHB) of protein extracts from three different embryonic time points: t1 (0–2 h), t2 (2–4 h), and t3 (4–6 h). This blot shows detectable levels of increasing NICD beginning at blastoderm cellularization. N-FL is full-length Notch. **(B)** Western blot using same NICD-specific antibody plus a *β*-tubulin antibody for sample loading. Lower levels of NICD are detected for the *opa35*-encoded (RALX237-bg1) N versus the *opa31*-encoded N (RALX822-bg1). The lanes for time point t2 (blank lines) were removed for increased clarity because sample quantity was low from both samples. Lower levels of NICD in lines carrying extreme *opa* length variants are consistent with observed phenotypic outcomes as explained below. **(C)** *AS-C* circuit implementing lateral inhibition via N-Delta signaling. The *AS-C*, *E(spl)-C*, and proneural genes are depicted in abstract gene logic diagrams with positive *cis*-regulatory inputs represented by receptor symbols and negative *cis*-regulatory inputs represented by repression symbols. These circuits stipulate that expression of *AS-C* is negatively correlated to NICD levels via its induction of the *E(spl)-C*. See text for details. **(D)** The transcriptional state space map is conceptually depicted so that transcriptional states are compressed into the *x*-*z* plane while the *y*-axis is reserved for depicting expression levels of *AS-C* genes. This conceptual map is intended to focus on two major points. The first point is that distinct pre-patterning signals will create different expression states for ectodermal cells (depicted are three such spaces of states, a–c). The second point is that dosage compensation at relevant X-linked loci in males may not be fully equivalent to female expression levels, causing slightly shifted blue and pink state space regions. Both points together may explain macrochaete-specific and sex-specific effects of altered NICD levels. **(E)** An altered transcriptional state space map is shown. Natural selection would allow the evolution of canalized developmental pathways capable of working within all of the female sub-spaces as well as the mostly overlapping male dosage compensated sub-spaces. However, genetically-shifted boundaries in functional state space regions (dotted areas) for specific bristles could affect males and females to different extents. **(F)** As ectopic macrochaetes and empty sockets are more common than missing sensory organs or split bristles in adults carrying the extreme *N opa* alleles, we suggest that the common defect is reduced levels of NICD either through decreased processing and/or increased degradation.

First, the observed enrichment for *opa* variants in the radiation-resistant RAL lines, including RAL-237, could be due to the expression of variant pQ Notch isoforms. These variant N isoforms could misfold and cause ER stress during co-translation and N processing (Fortini 2009; Yamamoto *et al.* 2010), and lead to chronic induction of UPR-related cell survival factors (Nagelkerke *et al.* 2014). Increased levels of Notch degradation would be a side consequence of these misfolding issues.

Second, if variant *opa* repeat lengths lead to reduced levels of Notch overall, this also would be consistent with increased levels of *AS-C* genes during macrochaete specification in the larval imaginal discs (Fig. 10C). Reduced levels of E(spl)-C bHLH repressors would be expressed in proneural cells, leading to increased expression of *AS-C*, and elevated rates of ectopic SOP specification (Fig. 10D, E). Similarly, during SOP lineage specifications, we would expect to see doubled sockets more frequently than doubled (*i.e.*, split) bristles emanating from a macrochaete position. As both ectopic bristles (doubled SOPs) and empty sockets are more frequently observed than missing (socket-less) bristles and split bristles, respectively, we suspect reduced NICD levels are behind *opa* variant phenotypes (Fig. 10F). The effects on specific bristles or the bristles of a specific sex (male or female) can also be explained by the different levels of *AS-C* and other transcriptional state space parameters expected for different proneural clusters and/or dosage compensation states.

## Discussion

We discovered a plethora of cryptic *N opa* variants by sampling the DGRP lines and genotyping multiple independent clones from each line by Sanger sequencing. We find that 47.4% of RAL haplotypes (18.5/39) do not correspond to the standard *opa31* length, and are not being accurately genotyped by high-throughput sequencing and assembly. Importantly, alleles characterized by extremely short or long *opa* repeats have common developmental defects, including classic *N* phenotypes affecting macrochaete patterning and wing notching. These phenotypes intensify when their X chromosomes or regions of the X chromosome containing *N* are outcrossed and/or recombined out into other backgrounds. The outcrossing effect suggests that these particular isogenic RAL lines contain suppressor modifiers. It also raises the question as to whether the 205 DGRP isofemale lines, established after 20 generations of inbreeding of the 1,500 Raleigh females, are enriched for compatible alleles (David *et al.* 2005). In any event, our results profoundly underscore the importance of developing new strategies for *de novo* whole-genome sequencing and assembly (Hahn *et al.* 2014). These strategies need to be open (verifiable) and ideally also suited to extension and customization.

Among the new *opa* variants, we discovered a rare short *opa23* allele that causes embryonic lethality in many backgrounds. This chromosomal region around the *N* locus is also responsible for producing notched wings when recombined into different backgrounds. Of all the rare *opa* alleles that we discovered by sampling DGRP RAL lines and other stocks, the *opa23* allele has the greatest length change relative to the wild-type *opa31*. It is also the only *opa* allele not encoding the histidine residue internal to the pQ tract.

Tautz reported hypervariability for the *N opa* repeats of *D. melanogaster* after seeing four length variants from 11 independent isogenic female lines established from North and South America, Europe, Asia, and Australia (Tautz 1989). These four alleles encode the pQ range of Q_13_HQ_17–20_ and predominantly correspond to wild-type *opa31* and *opa32* (81.8%). Two rare variant haplotypes corresponded to an *opa33c* encoding Q_13_HQ_19_, and an *opa34b* encoding Q_13_HQ_20_ (Fig. 2). If we consider just the three North American isofemale lines genotyped by Tautz, then these give a 2-to-1 ratio of *opa31* to *opa32*, which roughly approximates the 2.7-to-1 ratio we obtained from genotyping the isofemale Raleigh lines of the DGRP (Table 4). However, this ratio falls to 0.8-to-1 when all the world-wide isofemale lines are considered. Indeed, the single isofemale line from Japan was homozygous for *opa34b*, while the single isofemale line from the USSR was heterozygous for *opa33c*/*opa31*. Both of these variants have not yet been genotyped by us in the re-sequenced DGRP lines. Thus, it will be interesting to see if different *opa* alleles are found in different world populations.

**Table 4.** Comparison with world-wide distribution of Notch opa repeat variants

| Study | Source | # lines<sup>a</sup> | *opa*31 | *opa*32 | *opa*33 | *opa*34 | All other *opa* length variants |
| --- | --- | --- | --- | --- | --- | --- | --- |
| Rice et al. (this study) | North America | 40 | 51.2% | 18.7% | 12.5% | 2.5% | 12.5% |
| Tautz (1989) subset<sup>b</sup> | North America | 3 | 66.7% | 33.3% | 0.0% | 0.0% | 0.0% |
| Tautz (1989) all | world-wide | 11 | 36.4% | 45.4% | 4.5% | 9.1% | 0.0% |

<sup>a</sup> Number of iso-female lines genotyped
<sup>b</sup> 3/11 iso-female lines from world-wide populations (Tautz 1989)

The distribution of *opa* length variants suggests extreme purifying selection against the shorter would-be alleles in the barrens range (*opa* 24–30), and somewhat more moderate purifying selection against the many different longer variants in the *opa* allelic jungle range (*opa* 31–38). Given all the known *opa* genotypes (Fig. 2) and the *opa* variant phenotypic effects, the allelic barrens is unlikely to be an artifact of poor sampling. We propose that the existence of *opa23* is supportive of the reality of the barrens because it is missing the codon for histidine. We hypothesize that on the far short side of the barrens *opa* alleles may become semi-viable, provided the interrupting histidine is missing. In the presence of an intervening histidine residue, the *opa*-encoded pQ peptide may behave differently as seen in other contexts (Sharma *et al.* 1999; Sen *et al.* 2003; Kim 2013). In this regard it is interesting that the longest *opa* variant we have observed also has a contiguous stretch of 23 Q’s. Thus, 23 Q’s are the longest uninterrupted tracts we have observed regardless of whether we are talking about the short or long *opa* variants. Why this should be so must be specific to the roles played by NICD.

Perutz explained the 40 glutamine pathological threshold via the amyloid structure that is able to form from two stacked 20 residue-long rings (Perutz *et al.* 2002). In this cross-*β* amyloid nanotube each glutamine side group forms a hydrogen bond with another side-chain in the adjacent ring. These side-chain hydrogen bonds are in addition to the ones along the peptide backbone, typical of all *β*-sheet structures. However, the observed *opa* repeat distribution may also be indicating lethality of expansions past Q_23_. Because this is significantly below the pathological limit, we think either of two possible explanations will eventually be found to apply. First, a long pQ tract with only a single intervening histidine might behave similarly to an uninterrupted tract in some respects, such that a new Perutz amyloid threshold exists near 38–40 residues. Second, some other non-amyloid structure, possibly an interdigitated *β*-sheet interaction between NICD and one or more other interacting proteins exist. These interactions may be too strong when the tract is abnormally expanded. Similarly, the allelic barrens might represent the lethality of contracting the pQ tracts below Q_13_ (left-side) or Q_17_ (right-side) if doing so impacts specific critical interactions.

We find only three synonymous A/G (puRine) substitutions in all of the known *opa* sequences (Fig. 3). These three synonymous polymorphisms occur at specific positions within and near the long (CAG)_7_ triplet repeats (an unspecified number of the 11 Tautz sequences have the *2 polymorphism as it occurs in that consensus). The location of these synonymous substitutions is consistent with two empirical observations. First, indel mutations scale as a function of repeat number for diverse repeat unit lengths and sequences (Ananda *et al.* 2013). Second, nucleotide substitution rates tend to be elevated around indel polymorphisms (Ellegren 2000; Schlötterer 2000). At a superficial glance, the occurrence of these substitutions across the diversity of *opa* genotypes (Fig. 3) suggests they are independently recurrent mutations. However, we strongly caution against such a premature conclusion because the frequency of partial *opa* gene conversion events at this specific region is not known.

We also found several interesting correlations harbored by RAL lines carrying distinct *opa* variants and other loci encoding variant pQ-rich transcription factors (TFs). These include the TFs encoded by the *In(1)3R* inversion on chromosome 3R, and the Dorsal TF encoded on chromosome 2L. Previous work has already shown substantial variation and natural selection for the lengths of pQ tracts in the Notch co-activator Mastermind (Newfeld *et al.* 1991, 1993, 1994).

The striking correlation between cryptic yet functionally interacting repeats at unlinked loci suggests that key repeat variables underlie routine gene regulatory network adjustments. This is grave cause for concern given our facile focus on simple coding substitutions at both *cis*-regulatory and protein-coding DNA sequences. Furthermore, given the unusually high hydrogen bonding capacity for a Gln residue located inside a *β*-sheet structure (up to three H-bonds per Gln), variant pQ alleles are likely to contribute to developmental temperature norms as well as latitudinal and climatic population differentiation (David *et al.* 1997). Thus, during generation-to-generation selective adjudication of individual instances of a gene regulatory network, micro-satellite repeat variants likely rule the day, much like slow weathering and erosion are powerful geological processes that sculpt the Earth.

## Acknowledgments

This work was funded in part by an NSF CAREER award to A.E. (IOS:1239673) and a 2014 Evelyn Hart Watson research fellowship to C.R. We would like to thank Pamela Geyer and Josep Comeron for many helpful consultations.

